# Pre-assembled Cas9 ribonucleoprotein-mediated gene deletion identifies the carbon catabolite repressor and its target genes in *Coprinopsis cinerea*

**DOI:** 10.1101/2022.06.07.495237

**Authors:** Manish Pareek, Botond Hegedüs, Zhihao Hou, Árpád Csernetics, Hongli Wu, Máté Virágh, Neha Sahu, Xiao-Bin Liu, László G. Nagy

**Affiliations:** Institute of Biochemistry, Biological Research Centre, Temesvári krt. 62, 6726 Szeged, Hungary

**Keywords:** Carbon catabolite repression, Genome editing, Mushroom-forming fungi, CAZymes, Transcription factors (TFs), Split-marker, Transporters.

## Abstract

Cre1 is an important transcription factor that regulates carbon catabolite repression (CCR) and is widely conserved across fungi. This gene has been extensively studied in several Ascomycota species, whereas its role in gene expression regulation in the Basidiomycota remains poorly understood. Here, we identified and investigated the role of *cre1* in *Coprinopsis cinerea*, a basidiomycete model mushroom that can efficiently degrade lignocellulosic plant wastes. We used a rapid and efficient gene deletion approach based on PCR-amplified split-marker DNA cassettes together with *in-vitro* assembled Cas9-guide RNA ribonucleoproteins (Cas9-RNPs) to generate *C. cinerea cre1* gene deletion strains. Gene expression profiling of two independent *C. cinerea cre1* mutants showed significant deregulation of carbohydrate metabolism, plant cell wall degrading enzymes (PCWDEs), plasma membrane transporter-related and several transcription factor encoding genes, among others. Our results support the notion that, similarly to reports in the ascomycetes, Cre1 of *C. cinerea* orchestrates CCR through a combined regulation of diverse genes, including PCWDEs, transcription factors that positively regulate PCWDEs and membrane transporters which could import simple sugars that can induce the expression of PWCDEs. Somewhat paradoxically, though in accordance with other Agaricomycetes, genes related to lignin degradation were mostly downregulated in *cre1* mutants, indicating they fall under different regulation than other PCWDEs. The gene deletion approach and the data presented in this paper expand our knowledge of CCR in the Basidiomycota and provide functional hypotheses on genes related to plant biomass degradation.

**Importance:** Mushroom-forming fungi include some of the most efficient degraders of lignocellulosic plant biomass. They degrade dead plant materials by a battery of lignin-, cellulose-, hemicellulose- and pectin-degrading enzymes, the encoding genes of which are under tight transcriptional control. One of the highest-level regulation of these metabolic enzymes is known as carbon catabolite repression, which is orchestrates by the transcription factor Cre1, and ensures that costly lignocellulose-degrading enzyme genes are expressed only when simple carbon sources (e.g. glucose) are not available. Here, we identified the Cre1 ortholog in a litter-decomposer Agaricomycete, *Coprinopsis cinerea,* knocked it out and characterized transcriptional changes in the mutants. We identified several dozen lignocellulolytic enzyme genes as well as membrane transporters and other transcription factors as putative target genes. These results extend knowledge on carbon catabolite repression to litter decomposer Basidiomycota.

## Introduction

Carbon catabolite repression (CCR) is a gene regulatory mechanism that inhibits the utilisation of complex carbon sources in the presence of preferred carbon sources, such as glucose (Görke and Stülke, 2008; Adnan *et al*., 2017). The transcription factor Cre1 and its orthologs have been identified as key regulators of CCR and are extensively studied in several model fungal species, such as *Aspergillus* spp. (CreA), *Saccharomyces cerevisiae* (Mig1), and *Neurospora crassa* (Cre-1) (Fasoyin *et al*., 2018; Santangelo, 2006; Sun and Glass, 2011). In the presence of glucose, Cre-1 binds promoters and represses the transcription of genes responsible for utilisation of alternate carbon sources (Gancedo, 1998; Dowzer and Kelly, 1991; Ries *et al*., 2016). The absence of the preferred carbon source can lead to phosphorylation of CreA and a change in its subcellular localization, ultimately leading to derepression of its target genes involved in the utilisation of non-preferred carbon sources such as lignocellulosic plant remnants (Strauss *et al*., 1999; Kunitake *et al*., 2019; Hu *et al*., 2020; De Assis *et al*., 2021). It has been shown that manipulating the carbon catabolite repressor has an impact beyond CCR, on amino acid metabolism, intracellular trehalose levels or stress, among others (De Assis *et al*., 2021) suggesting that Cre1/CreA/CRE-1 can directly or indirectly regulate a large suite of metabolic and other genes.

Carbon catabolite repression is best-studied in the Ascomycota (De Vries and Makela, 2020, Adnan *et al*., 2017, De Assis *et al.,* 2021), whereas in the Basidiomycota several details of the process are poorly known, including upstream regulators and downstream target genes of Cre1. CCR has been demonstrated in several wood-decay fungi, including the white rotters *Pleurotus ostreatus* (Yoav *et al*., 2018; Alfaro *et al*., 2020)*, Dichomitus squalens* (Daly *et al*., 2019)*, Ganoderma lucidum* (Hu *et al*., 2020), *Phanerochaete chrysosporium* (Suzuki *et al*., 2008) and the brown rot fungus *Rhodonia placenta* (Zhang *et al*., 2022). This suggests that orthologs of Cre-1 should exist in the Basidiomycota as well. Indeed, deletion and overexpression of the *cre1* gene in the *P. ostreatus* showed that *cre1*-mediated regulation of CAZyme activities is substrate-dependent (Yoav *et al*., 2018). A recent study based on RNAi silencing showed that the creA ortholog in *G. lucidum* can regulate cellulase enzyme activity and transcription of related genes (Hu *et al*., 2020). However, despite these studies, the role of the *cre1* gene in global gene regulation, and its regulon in mushroom-forming fungi, are unclear, hampering in-depth studies on the regulation of plant biomass degradation in the Basidiomycota.

Here, we investigated the effects of *cre1* gene deletion on the phenotype and global gene expression regulation in *Coprinopsis cinerea*. *C. cinerea*, also known as "Inky-cap mushroom’’ forms complex fruiting bodies that autolyse to produce a black mass of spores. It is a widely used model in mushroom developmental biology (Kües, 2000; Pukkila, 2011). At the same time, *C. cinerea* is a litter decomposer species, which is adapted to decomposing lignocellulosic substrates such as manure, hay or straw. Litter decomposers represent an ecologically successful guild among mushroom-forming fungi, mostly in the Agaricales (Floudas *et al*., 2020; Ruiz-Duenas *et al*., 2021). Unlike white and brown rot fungi which decompose wood, litter decomposers obtain carbon from non-woody plant biomass. In *C. cinerea*, the availability of a self-fertile homokaryotic strain (AmutBmut; *pab1-1*), along with widely used selection markers, including auxotrophic marker genes, facilitate genetic studies (Swamy *et al*., 1984, Dörnte *et al*., 2020, Kües, 2000).

Genomic integration of foreign DNA occurs either by non-homologous end joining (NHEJ) or homologous recombination (HR). NHEJ is widespread in fungi, while HR is very limited (Krappmann, 2007; Ishibashi *et al*., 2006), especially in the mushroom-forming fungi (Ohm *et al*., 2010; Salame *et al*., 2012), which complicates HR-dependent targeted gene deletion. The ratio of HR-mutants has been increased by disruption of the NHEJ genes *ku70*, *ku80* or *lig4* in mushroom-forming fungi (Ninomiya *et al*., 2004, Ishibashi *et al*., 2006, Nakazawa *et al*., 2011, De Jong *et al*., 2010). Another approach for preferentially utilising HR dependent gene integration is the split-marker gene deletion approach, which uses two overlapping fragments of the dominant marker with flanking target gene fragments (Chung and Lee, 2015). In *Cordyceps militaris*, the split-marker approach results in increased gene deletion compared to a single fragment (Lou *et al*., 2018). This approach is commonly used for genetic manipulations in filamentous fungi (Kück and Hoff, 2010), but not in *C. cinerea*. Recently, genome editing using Clustered Regularly Interspaced Short Palindromic Repeats (CRISPR)/CRISPR-associated protein 9 (Cas9) has become popular in various fungi including Agaricomycetes (Schuster and Kahmann, 2019, DiCarlo *et al*., 2013; Raschmanová *et al*., 2018, Wang and Coleman, 2019; Sugano *et al*., 2017; Vonk *et al*., 2019; Liu *et al*., 2020; Boontawon *et al*., 2021a) and involves a report of using the split-marker together with the pre-assembled Cas9 ribonucleoproteins (Cas9-RNPs) in *P. ostreatus* (Boontawon *et al*., 2021b).

In this study, we identified the *C. cinerea cre1* gene and generated two independent *C. cinerea cre1* mutants using pre-assembled Cas9-RNPs along with a PCR-based split-marker DNA repair cassette. Transcriptional changes were analysed in the two *C. cinerea cre1* mutants using Quant-Seq. Our analysis highlights putative target genes of Cre1 in a litter decomposer basidiomycete. We detected differential expression of several carbohydrate metabolism genes, carbohydrate-active enzyme encoding genes (CAZymes) related to plant cell wall degradation, plasma membrane transporters, and several transcription factors. In addition, this work also provides a simple, efficient and rapid approach for gene deletion in the mushroom *C. cinerea*.

## Materials and Methods

### Strain, culture conditions, media, buffers and reagent preparation

The homokaryotic strain *Coprinopsis cinerea* AmutBmut; *pab1-1* (FGSC 25122) was used for the study (Swamy *et al*., 1984). The fungus was cultured on YMG medium at 37°C under continuous light for 6-7 days to obtain oidia for protoplast preparation (Xie *et al*., 2020). The compositions and preparations of YMG, minimal medium, regeneration medium, top agar, MM buffer, MMC buffer, and PEG /CaCl_2_ followed recipes described by Dörnte and Kües, 2012.

### Comparative genomics and phylogenetic analysis

The Cre1 (hereafter, "Cre1’’ indicates protein and "*cre1*" indicates gene) ortholog (JGI protein ID: 466792) was identified in *C. cinerea* AmutBmut based on reciprocal best blast hits using the *Neurospora crassa* Cre-1 (NCU08807) protein as query. The nucleotide sequence was retrieved from MycoCosm maintained by JGI for *Coprinopsis cinerea* AmutBmut https://mycocosm.jgi.doe.gov/Copci_AmutBmut1/Copci_AmutBmut1.home.html (Muraguchi *et al*., 2015).

A Maximum Likelihood phylogenetic analysis was performed to confirm orthology of Cre1 to carbon catabolite repressors from Ascomycota. For this purpose, we selected orthologs represented by reciprocal blast hits for key Basidiomycota (*C. cinerea*, *D. squalens*, *Laccaria bicolor*, *Lentinula edodes*, *P. ostreatus*, *Schizophyllum commune*, *Trametes versicolor*, and *Ustilago maydis*) and Ascomycota (*Aspergillus nidulans*, *Aspergillus niger*, *N. crassa*, *S. cerevisiae*, *Trichoderma reesei*, and *Yarrowia lipolytica*). For rooting the tree, we chose sequences of related C_2_H_2_ transcription factors of *S. cerevisiae* and *Aspergillus* spp. Protein sequences were retrieved from the MycoCosm or Uniprot databases or from the Saccharomyces Genome Database (Cherry *et al*., 2012), aligned with PRANK version 170427 (Löytynoja, 2014), and trimmed using the built-in ‘strict’ settings of trimAl (Capella-Gutiérrez *et al*., 2009). Accession numbers are provided in **Fig. 1**. The resulting multiple sequence alignment was analysed using RAxML version 7.0.2 under the PROTGAMMAWAG model of evolution with 1,000 rapid bootstrap replicates to assess branch support. Trees were rooted and visualized in FigTree v1.4.4 (Rambaut, 2018).

**Fig. 1.**
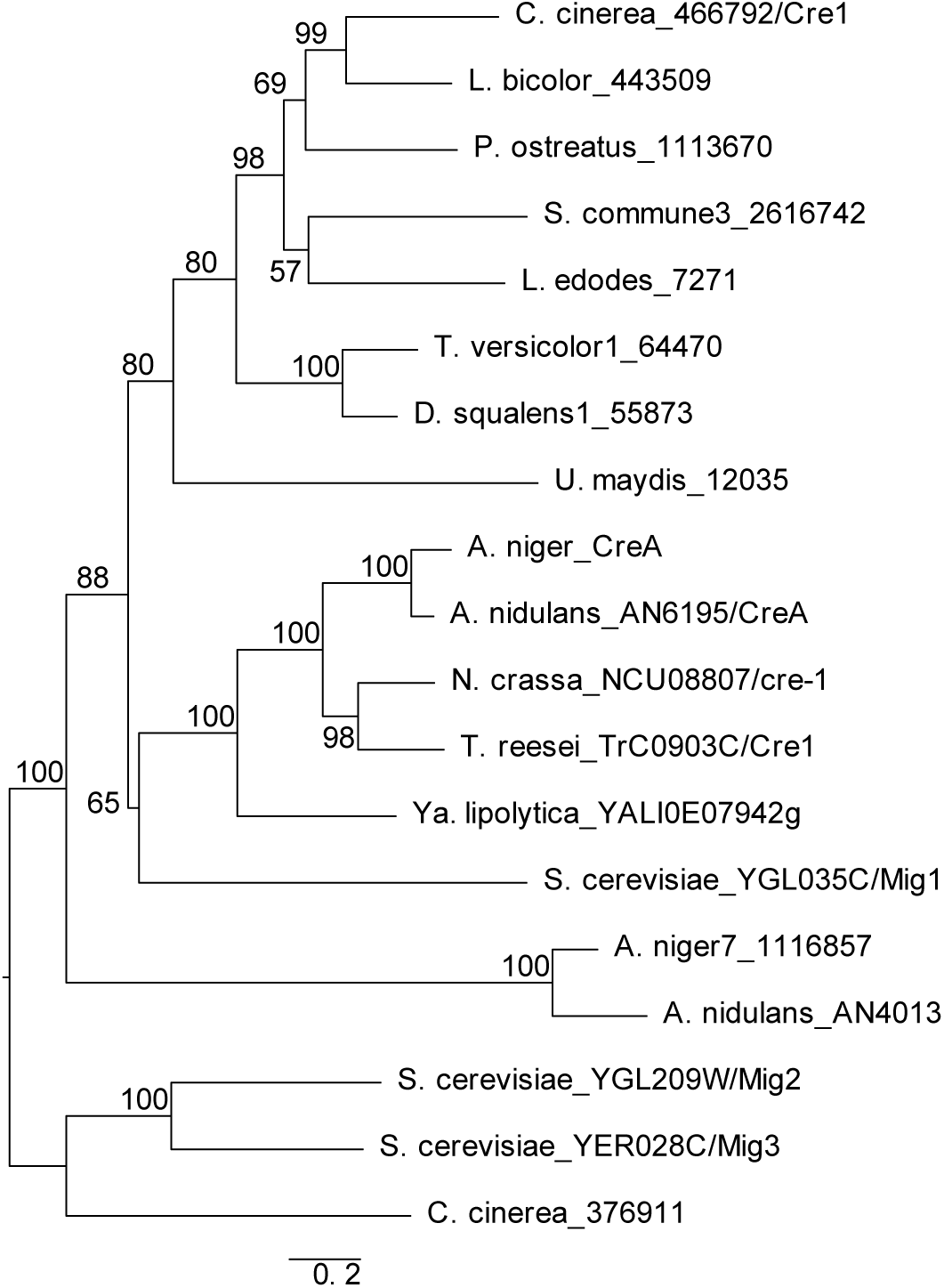
Maximum likelihood phylogenetic tree of C_2_H_2_ zinc finger transcription factors of selected filamentous fungi and yeasts. The tree reveals the monophyly of *C. cinerea* Cre1 (466792/Cre1) with experimentally characterised members of the family including *S. cerevisiae* Mig1, *A. niger* CreA*, A. nidulans* CreA, *N. crassa* Cre-1 and of *T. reesei* Cre1. The numbers next to branches represent the bootstrap support values. JGI protein IDs are given for each protein used in the analysis.

### sgRNAs design and in-vitro assembled Cas9-RNPs preparation

The tracrRNA and crRNA were commercially synthesised for the *in-vitro* assembled RNPs (IDT, USA). sgRNAs protospacer were designed with the sgRNACas9 software using the *C. cinerea cre1* gene sequence (protein ID: 466792) (Xie *et al*., 2014). Two sgRNAs (*cre1_exon_S_8* and *cre1_exon_S_156*) from the 5’ and 3’ ends of the *cre1* gene were selected which has minimal off-target effects in the *C. cinerea* genome (**Table S1**). The position of the sgRNAs binding on the *C. cinerea cre1* gene locus is shown in **Fig. S1 A**. Each synthesised crRNA and tracrRNA was annealed and mixed with the commercially available Alt-R® S.p.Cas9 Nuclease V3 (IDT, USA) along with Cas9 buffer to obtain the *in-vitro* assembled RNP complexes according to the manufacturer’s instructions. Briefly, for each *in-vitro* assembled RNP mixture, 12 μl of equimolar RNA duplexes were first formed by mixing 1.2 μl of crRNA with 1.2 μl of tracrRNA from the stocks of crRNA (100 μM) and tracrRNA (100 μM) together with 9.6 μl of duplex buffer (IDT) and then incubated at 95°C for 5 minutes and cooled for 2 minutes. Next, 12 μl RNA duplex, 0.5 μl Cas9 (10 μg/μl), 1.5 μl Cas9 working buffer, and 1.0 μl duplex buffer were mixed and incubated at 37°C for 15 minutes. Fifteen μl of RNP mix was used for protoplast transformation. One RNP reaction for each *cre1_exon_S_8* and *cre1_exon_S_156* sgRNA was prepared and used along with the purified split marker DNA repair cassette for protoplast transformation.

### PCR based split-marker DNA repair cassette preparation

Para-aminobenzoic acid (PABA) auxotrophy in the *C. cinerea* AmutBmut; *pab1-1* strain can be used for the selection of the transformants with *pab1* gene as selectable marker (Bottoli *et al*., 1999). The split-marker DNA cassettes for the *pab1* (PABA) marker gene along with ∼700 bp upstream homology arm (UHA) and downstream homology arm (DHA) of *C. cinerea* were prepared using the double-joint PCR method (DJ-PCR) (Yu *et al*., 2004). First, 1 kb upstream of the 5’ end (UHA) and 1kb downstream of the 3’ end (DHA) were amplified with the primers (P1, P2 and P3, P4, respectively). The *pab1* cassette along with the promoter and terminator (CcPAB1, 3097 bp) was amplified with primers P5 and P6 from vector pMA412 and used as a selection marker for *C. cinerea* (AmutBmut; *pab1-1*) (Stanley, 2014, Schmieder *et al*., 2019). P5 and P6 primers have overlapping overhangs with *C. cinerea cre1* UHA and DHA fragments. The vector map for the split marker cassettes (S1 and S2) along with primer locations can be found in **Fig. S1 B**. Each purified 1kb UHA or DHA fragment was fused to the 3kb PABA cassette in separate tubes to generate a long common DNA fragment using the DJ-PCR protocol (UP-PABA or PABA-DOWN) (Yu *et al*., 2004) with Phusion Green Hot Start II High-Fidelity PCR Master Mix (Thermo Scientific™). Finally, Split1 (S1) cassette was amplified from UP-PABA using P7 and P8. Similarly, Split 2 (S2) was amplified from the PABA-DOWN using P9 and P10. Both S1 and S2 have ∼1081 bp long overlapping *pab1* marker gene fragments, which is required for recombination between split DNA fragments. The protocol for generating a split-marker DNA repair cassette for gene deletion is described in more detail elsewhere (Catlett *et al*., 2003, Yu *et al*., 2004, Pareek *et al*., 2019). All primer positions are shown in **Fig. S1 A and B** and their DNA sequences are given in **Table S1**. Primers for the DJ-PCR were designed using the Gibson assembly primer design function of SnapGene software (from Insightful Science; available at snapgene.com). **Fig. S2** shows the purified S1 and S2 DNA cassette used for protoplast transformation.

### PEG-mediated protoplast transformation using Cas9-RNPs and split-marker DNA cassette mix

*In-vitro* assembled Cas9-RNPs for both ends of the *C. cinerea cre1* gene were used together with 10 μg of the amplified split-marker DNA (S1 and S2) for PEG mediated transformation of *C. cinerea* protoplasts as described by Dörnte and Kües (2012) with minor modifications. Briefly, 0.5 x 10^8^ oidia were harvested and washed with MM buffer, then resuspended in 950 μl MM buffer and mixed with 50 μl Protoplast F enzyme (Megazyme, Ireland). The mixture was incubated for at least 2.5 - 3 hrs to generate ∼70-90% protoplast in suspension. The protoplast suspension was washed with MMC buffer to stop the reaction and then resuspended in 200 μl of MMC buffer for two transformation reactions. Further, the protoplast solution was divided into two Falcon tubes, one for the split-marker DNA repair cassette alone transformation, and another for the RNPs together with the split-marker DNA repair cassette. To one tube 25 μl PEG /CaCl_2_, each RNP solution (15 μl), and 10 μg of the split-marker DNA repair cassettes (S1 and S2), and to another tube similar components except the RNP solution were added. The putative fungal transformants were picked from the minimal medium selection plates between 3-6 days of incubation at 37°C. Each colony was separately picked and placed on separate minimal medium plates. Genomic DNA was then isolated and selected fungal transformants were further analysed using PCR.

### PCR screening of the transformed colonies for C. cinerea cre1 gene deletion

Colony PCR was performed using the crude genomic preparation protocol described by Dörnte and Kües, (2013). Three screening methods were used for PCR screening and confirmation by sequencing (**Fig. S3**). Screening method 1: Primers P11 and P12 provide a size difference between the *C. cinerea cre1* wild-type allele (3.36 Kb) and the *pab1* cassette (3.62 Kb) in the putative transformants. All colonies were screened again using screening method 2, with one primer (P13) upstream of the UHA and another primer (P14) located within the *C. cinerea cre1* gene locus. We used 34-35 PCR cycles to identify the true deletion strain with complete removal of the *cre1* gene locus. All mixed and ectopic transformants should yield a PCR signal in screening method 2, but a true *cre1* gene deletion strain does not yield an amplicon.

For screening method 3, selected colonies were grown again on YMG medium and then genomic DNA was isolated from the fungal mycelium using the CTAB method (Zhang *et al*., 1996). PCR was used to amplify the large 3.0 kb upstream fragments (UP) using the external forward primer (P13) and the PABA internal primer (P8). Similarly, the PABA internal forward primer (P9) was used together with the external downstream reverse primer (P15) for amplification of the downstream fragment (DW). These amplified PCR fragments indicate the integration of the *pab1* marker into the *cre1* locus of *C. cinerea*. Finally, the UP and DW fragments were used for DNA sequencing with primers P13 and P16 (UP) and P15 and P17 (DW). All primer positions and DNA sequences are shown in **Fig. S1** and **Fig. S3** and **Table S1**. Later, two internal primers for *C. cinerea cre1* gene (P18 and P19) were also used for PCR confirmation of mutants from genomic DNA (**Fig. S1 A**).

### RNA isolation and sequencing

RNA was isolated from wild type (*Coprinopsis cinerea* AmutBmut; *pab1-1*) strain (3 replicates), the Δ*cre1*-43 and Δ*cre1*-91 mutants (2 replicates for each independent mutant). All colonies were grown on YMG medium (0.2% glucose) overlaid with cellophane for 84 hrs in the dark at 28°C. Mycelia were collected and homogenised in liquid nitrogen using a mortar and pestle. Total RNA was isolated using the Quick RNA mini kit (Zymo Research, USA). Preparation of cDNA libraries for Quant-Seq, purification, and sequencing were performed by iBioScience Ltd (Pécs, Hungary) on an Illumina NextSeq instrument to achieve a sequencing depth of at least 10 million reads per sample.

### Quant-Seq data analysis

The reference genome sequence (*Coprinopsis cinerea* Amut1Bmut1 pab1-1 v1.0) and gene annotation (Copci_AmutBmut1_GeneModels_FrozenGeneCatalog_20160912) were downloaded from the JGI database. Read quality was checked using FastQC (Andrews, 2010). Adapter sequences, polyA reads, and low-quality tails were removed using bbduk version 38.92 (Bushnell *et al*., 2017). Trimmed reads were mapped to the reference genome using STAR version 2.6.1a_08-27 (Dobin *et al*., 2013). Trimming and mapping parameters were set according to the manufacturer’s recommendations (Lexogen, Austria). Only CDS gene annotations were considered in the read to gene assignment step with a 400 bp overhang added at the 3’ ends of each gene. For the assignment, the featureCounts version Rsubread_2.6.4 (Liao *et al*., 2014) function of the Rsubread package was used. For differential gene expression analysis, the edgeR version 3.34.0 (Robinson et al., 2010) and limma version 3.48.3 (Ritchie et al., 2015) packages were used. Genes with a fold change ≥ 2 and a Benjamini-Hochberg (BH) adjusted p-value ≤ 0.05 were considered upregulated (deg_status = 1). Genes with a fold change ≤ -2 and a Benjamini-Hochberg (BH) adjusted p-value ≤ 0.05 were considered downregulated (deg_status = -1). We also generated the list of IPR terms, IPR descriptions, and GO terms associated with the *C. cinerea* protein ids based on the output of InterPro Scan v86 (**Table S2**). We analysed the Quant-Seq data of Δ*cre1*-43 and Δ*cre1*-91 mutants independently to find differentially expressed genes compared with wild type samples, the data obtained are shown in **Table S3** and **Table S4**, respectively. The normalised average count per million (avg CPM) reads for the mutants and wild type is shown in **Table S5**. Raw reads corresponding to samples used in this study have been submitted to SRA under the accession number XXXXX (to be provided upon publication).

### Functional annotations

GO enrichment analyses were performed for differentially expressed genes separately for the Δ*cre1*-43 and Δ*cre1*-91 mutants using the topGo R package (Alexa and Rahnenfuhrer, 2021). We used the classicFisher algorithm to identify GO terms with a p-value ≤ 0.05. GO terms for all *C. cinerea* protein ids can be found in **Table S2**.

The putative CAZymes in *C. cinerea* were annotated along with their major substrate based on previous publications (Nagy *et al*., 2016, Strasser *et al*., 2015, Murphy *et al*., 2011, Floudas *et al*., 2012, Floudas *et al*., 2015, Kohler *et al*., 2015, Rytioja *et al*., 2014, Grunwald 2016, Miyauchi *et al*., 2017, Makinen *et al*., 2019, Krizsan *et al*., 2019, Miyauchi *et al*., 2020) and used for identification of plant cell wall degrading (PCWDE) and fungal cell wall (FCW) related CAZymes in our data. Putative transcription factors (TFs) were identified based on lists provided by Krizsan *et al*., (2019). Putative transporter proteins localised in the plasma membrane were taken from the list provided by Sahu *et al*., (2021). We also confirmed their subcellular localization with WoLF PSORT and used proteins with a predicted plasma membrane localization for analysis (Horton *et al*., 2006). Expression heatmaps were generated using the online tool HeatMapper (Babicki *et al*., 2016) and Venn diagrams were generated using InteractiVenn (Heberle *et al*., 2015).

### Fruiting assays

For analysis of fruiting body development, *C. cinerea cre1* mutants and the wild type strain (*Coprinopsis cinerea*, AmutBmut; *pab1-1*) were cultured on YMG for 5 days at 37°C in the dark and then transferred to a 12/12h light/dark cycle at 28°C for 5-7 days to analyse fruiting body development.

## Results

### Identification of the C. cinerea cre1 gene and phylogenetic analysis

We identified the Cre1 ortholog for *C. cinerea* using a reciprocal best hit strategy in protein BLAST and confirmed orthology using maximum likelihood analysis of the Cre1 orthologs and related C_2_H_2_ transcription factors (**Fig. 1**). The phylogeny strongly supported the monophyly of protein 466792 of *C. cinerea* with experimentally characterised members of the family, including Mig1 of *S. cerevisiae*, CreA of *Aspergillus niger* and *A. nidulans*, Cre-1 of *Neurospora crassa* and Cre1 of *Trichoderma reesei*. On the other hand, sequences representing Mig2 and Mig3 of *S. cerevisiae* and closely related proteins of *A. niger* and *A. nidulans* were clustered separately. Based on these results, we designate this protein of *C. cinerea* as Cre1.

### Preassembled Cas9-RNPs together with a split-marker DNA repair cassette increase the number of transformants

*C. cinerea* protoplast transformation using pre-assembled Cas9-RNPs together with a split-marker DNA repair cassette was compared with the split-marker DNA repair cassette alone method. We observed that the number of transformed colonies increased when RNPs were added together with the split-marker cassette compared with the split-marker DNA repair cassette alone transformation. Transformation with the split-marker DNA cassette alone yielded 78 colonies, whereas RNPs together with the split-marker DNA cassette in the same experiment yield 523 colonies within 5 days after transformation. Later, 96 colonies (48-48 from each method) were screened for deletion of the *C. cinerea cre1*.

### Identification of the C. cinerea cre1 gene deletion strains

We used the three PCR screening methods described in the experimental procedures to identify deletion strains among the putative transformants (**Fig. S3**). Using screening method 1, the replacement of target gene by the *pab1* marker cassette can be detected by the difference in size of the PCR amplicon compare to wild type fungal strain (**Fig. S4 A and B**). Screening method 2 is expected to show that true *cre1* deletion strains do not amplify the native *C. cinerea cre1* locus (**Fig. S4 C and D**). Considering the above two screening methods, three strains 43, 78 and 91 were selected for confirmation by DNA sequencing. All three strains were generated using the split-marker DNA repair cassette together with Cas9-RNPs. Final confirmation of the three deletion mutants by screening method 3 was performed by sequencing the upstream (UP) and downstream (DW) fragments as explained in the experimental procedures. The UP and DW DNA fragments amplified from all three deletion strains and their DNA sequencing results were aligned with the *C. cinerea cre1* gene locus or the PABA transformation cassette are shown in **Fig. 2 A-C**.

**Fig. 2.**
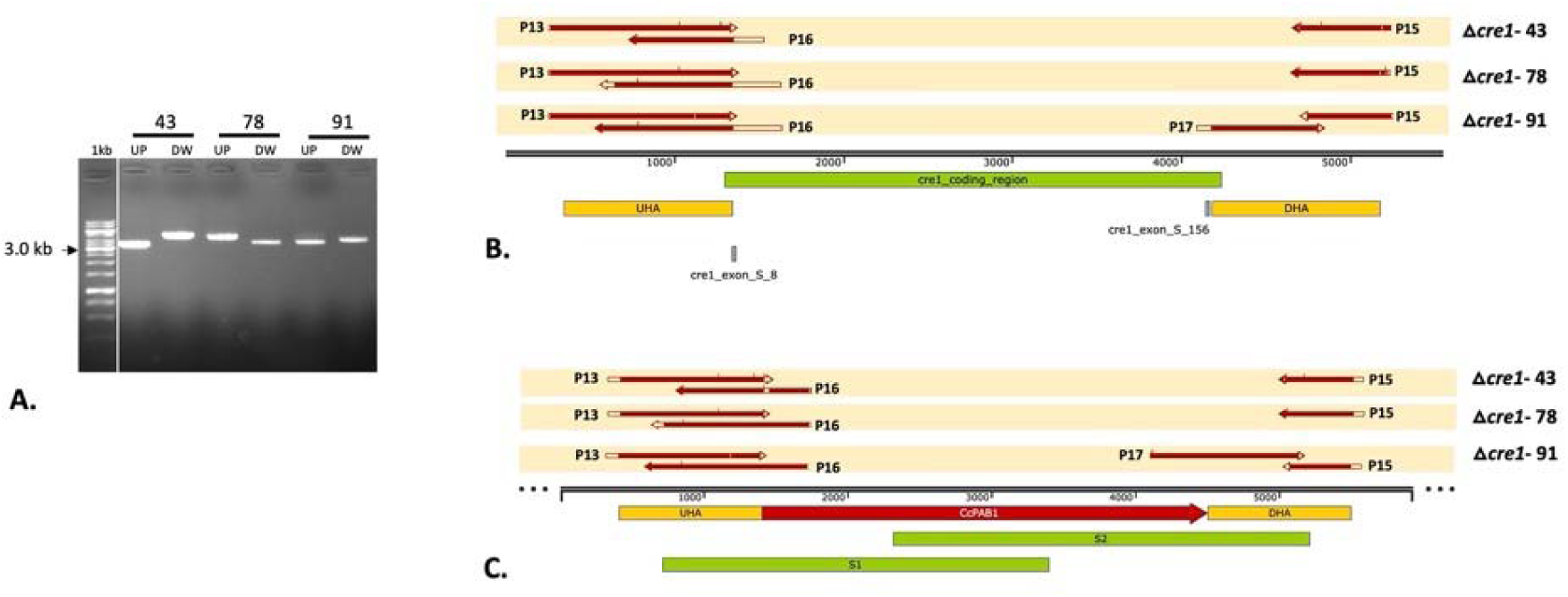
Confirmation of the *C. cinerea cre1* mutants by PCR amplification and DNA sequencing of *pab1* marker gene along with flanking region. PCR amplification of the upstream (UP) and downstream (DW) DNA fragments from the genomic DNA of the selected strains (43, 78 and 91) using the specific primers (A). The result of DNA sequencing of UP and DW PCR amplified fragments was aligned with the *C. cinerea cre1* gene locus (B) and the PABA transformation cassette (C). UHA is the upstream homology arm, DHA is the downstream homology arm and CcPAB1 is the *pab1* selection marker gene, cre1_coding_region is the *C. cinerea cre1* gene locus, cre1_exon_S_8 and cre1_exon_S_156 are the positions of the sgRNAs binding. UP and DW are the DNA fragments used for DNA sequencing with primers P13, P16, P17 and P18 as described in the experimental procedures.

The results of DNA sequencing of the UP fragment using the P13 primer for all three deletion strains Δ*cre1*-43, Δ*cre1*-78, and Δ*cre1*-91 were aligned to the UHA. It shows that the UP fragment amplified in the (**Fig. 2 A**) for all strains is from the *C. cinerea cre1* gene locus. Similarly, DNA sequencing reads of the UP fragment with the P16 primer aligned to the PABA gene but not to the *C. cinerea cre1* gene. This means that the PABA transformation cassette replaced the *C. cinerea cre1* gene locus in all three transformants. Similarly, the results of DNA sequencing of the DW fragment with the P15 primer aligned to DHA, confirming the position of the DW fragment at the *C. cinerea cre1* gene locus for all three deletion strains. DNA sequence data obtained with the P17 primer aligned to the PABA gene in the Δ*cre1*-91 deletion strain, but good sequencing results could not be obtained for the Δ*cre1*-43 and Δ*cre1*-78 mutants (**Fig. 2 B and C**). DNA sequencing of the UP and DW fragments amplified from the genomic DNA of the mutants and their alignment to the *cre1* PABA transformation cassette showed integration of the PABA gene into the *C. cinerea cre1* locus (**Fig. 2 B and C**). Our results show that all three selected strains (Δ*cre1*-43, Δ*cre1*-78, and Δ*cre1*-91) can be considered as *C. cinerea* mutants. Only mutants Δ*cre1*-43 and Δ*cre1*-91 were used for further analysis.

Although we performed very stringent PCR screening for the identification of the clean mutants, it was later found that the genomic DNA of both mutants gave PCR amplification of the small *C. cinerea cre1* gene fragment at a high number of PCR cycles using two gene internal primers (P18 and P19). Even after oidiation of the mutant strains, the PCR DNA band persisted. PCR amplification of the *C. cinerea cre1* fragment may be due to some cells having wild-type nuclei. This is not entirely clear and needs further investigation. Despite this uncertainty, we are confident that Δ*cre1*-43 and Δ*cre1*-91 represent *C. cinerea cre1* gene deletion mutants, which is confirmed by the results of Quant-Seq analysis (see below).

### C. cinerea cre1 knockout does not affect fruiting body formation

Both deletion strains Δ*cre1*-43 and Δ*cre1*-91 developed a fruiting body on YMG medium similar to the wild-type strain, no significant change was observed. (**Fig. 3**).

**Fig. 3.**
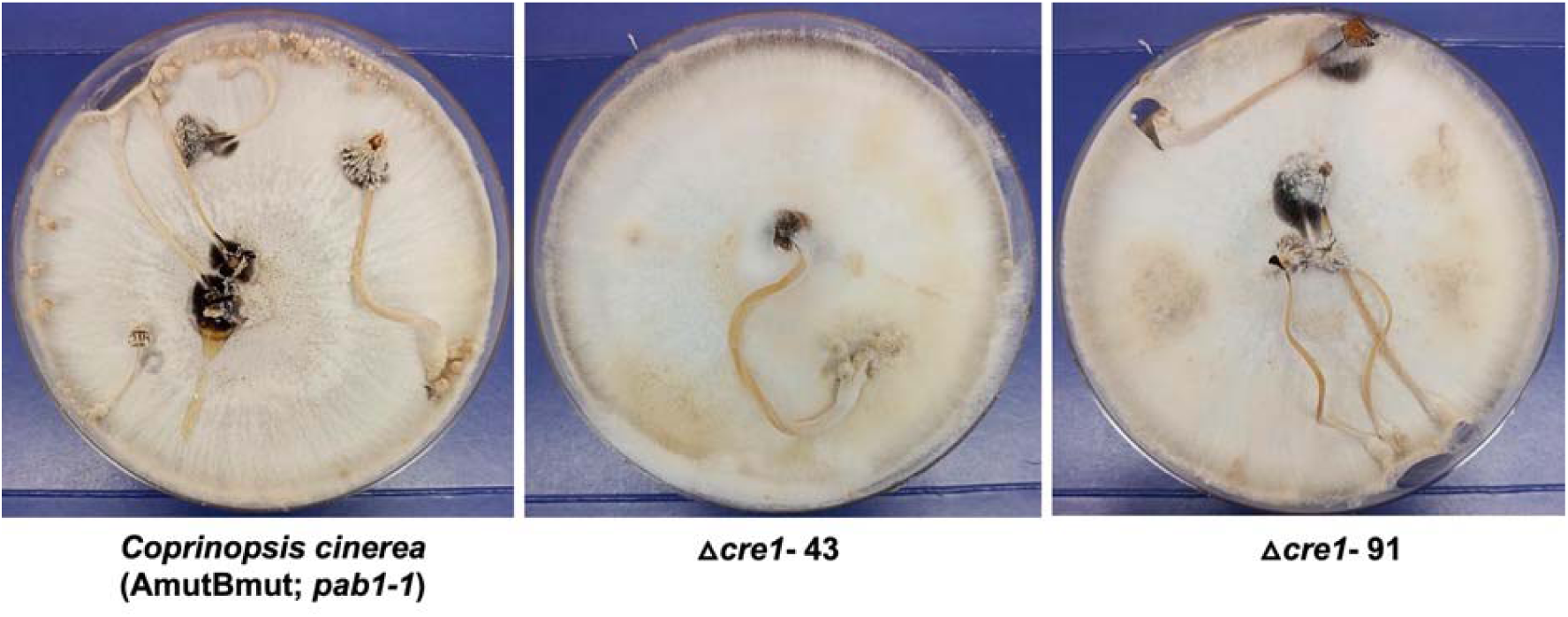
Fruiting body development in the wild type and *C. cinerea cre1* mutants. Development of fruiting bodies in *Coprinopsis cinerea* (AmutBmut; *pab1*) wild-type strain, Δ*cre1*-43 and Δ*cre1*-91 deletion strains after 14 days on YMG medium under standard conditions.

### Carbohydrate metabolism is broadly affected in cre1 mutants

In the Quant-Seq data for the *C. cinerea cre1* mutant and wild-type samples, approximately 74-94% of the reads were uniquely mapped to the *C. cinerea* genome (**Table S5**). We analysed the effects of *C. cinerea cre1* deletion on the transcriptome. The differentially expressed genes in both mutants are listed in **Table S3 and S4**. In the Δ*cre1*-43 and Δ*cre1*-91 mutants, 938 and 1,081 genes, respectively, were significantly differentially expressed (P≤0.05, |FC|>2) compared to wild type strains. The data show that 592 genes were common to both mutants. In the Δ*cre1*-43 mutant, 658 genes were upregulated and 280 genes were downregulated, whereas in the Δ*cre1*-91 mutant, 682 genes were upregulated and 399 genes were downregulated. Of these, 401 and 191 genes are commonly up and downregulated in Δ*cre1*-43 and Δ*cre1*-91, respectively.

Significantly enriched GO terms were identified for the differentially expressed genes separately for both mutants separately for the up- and down-regulated genes. The enriched GO terms largely overlapped between the two mutants, indicating similar transcriptomic changes after deletion of *C. cinerea cre1* (**Table S6**). Common enriched terms in the upregulated genes of *C. cinerea cre1* mutants showed a clear signal for processes related to carbohydrate metabolism (*e.g.* carbohydrate/organic/primary metabolic process, carbohydrate/cellulose/polysaccharide binding, hydrolase activity, extracellular region) (**Table S6)**. Interestingly, the upregulated genes of the *C. cinerea* Δ*cre1*-91 mutant were also enriched in terms related to transcription initiation and protein-DNA complex, which was not the case for the Δ*cre1*-43 mutant. GO terms related to arabinose metabolism and oxidoreductase activity were enriched only in the Δ*cre1*-43 mutant. Among the downregulated genes, oxidoreductase activity, iron binding, transmembrane transporter activity, extracellular space, and membrane-related terms were commonly enriched in both mutants. GO terms of interest related to G protein-coupled receptors (GPCR) can be observed in both mutants, whereas terms related to amino acid metabolism are mainly enriched in the Δ*cre1*-91 mutant (**Table S6**).

### Plant cell wall degrading CAZyme genes are broadly upregulated in cre1 mutants

We analysed the differentially expressed genes of Δ*cre1*-43 and Δ*cre1*-91 mutants for the presence of genes encoding carbohydrate-active enzymes (CAZymes). Of the 485 CAZymes in *C. cinerea*, we found that 101 and 102 were differentially expressed in the Δ*cre1*-43 and Δ*cre1*-91 mutant strains, respectively. Of these, 80 CAZymes were upregulated in each of the strains and 69 were common to both. In addition, 21 and 22 were downregulated in the Δ*cre1*-43 and Δ*cre1*-91 mutants, respectively, of which 13 were common. We classified CAZymes by their main substrate into groups acting on plant cell wall-degrading (PCWDE) and fungal cell wall (FCW) CAZymes following Sahu *et al*., (2021).

Of the differentially expressed genes in the *C. cinerea cre1* mutants, 82 genes were related to PCWDEs, 19 genes to FCW and 20 genes to neither PCWDE nor FCW. Overall, 35.49% (82/231) PCWDE genes, 17.92% FCW genes (19/106) and 12.82% (20/156) non-PCWDE and non-FCW genes were differentially expressed in the *C. cinerea cre1* mutants. This indicates that PWCDEs are enriched among the differentially expressed CAZymes.

We found a total of 67 upregulated PCWDE genes, of which 53 genes are commonly upregulated in both mutants (**Fig. 4A; Table S7**). In terms of predicted substrate, cellulase-encoding genes were dominant in both mutants, followed by hemicellulase and pectinase encoding genes (**Fig. 4A**). We found no gene predicted to be related to lignin degradation among the upregulated ones, though it should be noted that as a litter decomposer, *C. cinerea* possesses fewer lignin-related genes than white rot fungi (Floudas *et al*., 2020). Fifteen PCWDE genes were downregulated **(Fig. 4 A-B**; **Table S7**). These included four predicted lignin modifying enzyme genes (LMEs), one class-II peroxidase (AA2, protein ID: 446928) and three multicopper oxidases from the AA1 family (**Table S7**).

**Fig 4.**
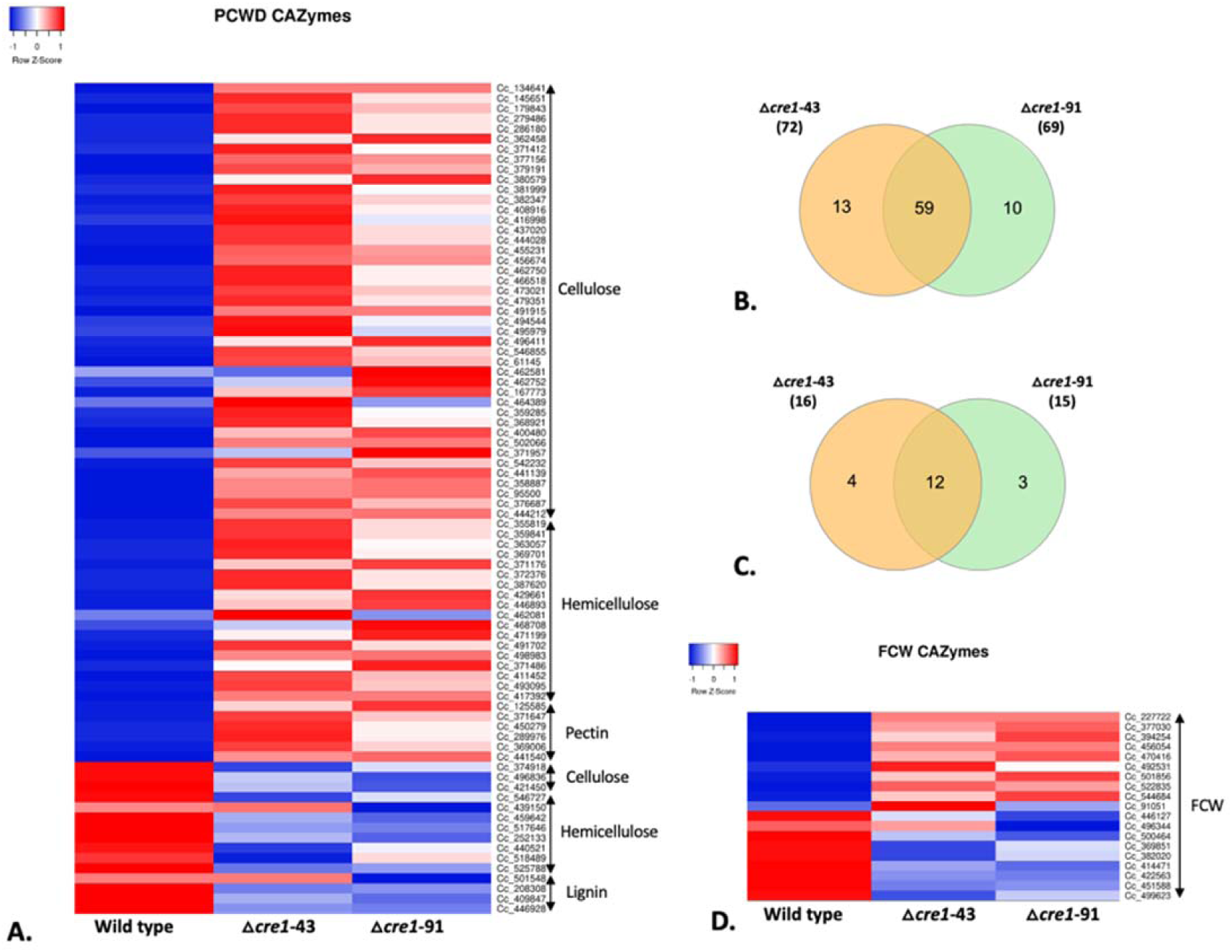
Differentially expressed CAZyme genes in the *C. cinerea* Δ*cre1*-43 and Δ*cre1*-91 mutants compared with wild type strain. (**A**) Heatmap showing the average CPM of differentially expressed plant cell wall degrading enzymes (PCWDEs) genes (JGI protein IDs) according to their main substrate of action (cellulose, hemicellulose, pectin and lignin) for the wild type, Δ*cre1*-43 and Δ*cre1*-91 mutants. (**B and C**) Venn diagram showing the common significantly differentially expressed PCWDE and FCW genes between the Δ*cre1*-43 and Δ*cre1*-91 mutants, respectively. (**D**) Heatmap showing the average CPM for the differentially expressed fungal cell wall (FCW) related CAZymes genes (JGI protein IDs) for the wild type, Δ*cre1*-43 and Δ*cre1*-91 mutants.

Among the differentially expressed cellulose-related CAZymes the Auxiliary Activity Family 9 (AA9) family and CBM1-domain containing genes were most abundant. Fifteen AA9 genes were differentially expressed, which are predicted to encode proteins with copper-dependent lytic polysaccharide monooxygenase (LPMOs) activity, and interestingly, all were upregulated in the *cre1* deletion strains except one. We also found 17 CBM1 domain containing genes among upregulated ones (none among downregulated); most of these were predicted to have enzymatic activities and belong to the GH5, GH10, GH11, GH30, GH62 and GH131 families. Other differentially expressed genes were found in GH families included GH1, GH3, GH5_22, GH6 and GH7. Two expansins were upregulated in the mutants, but two were also downregulated. Expansins have been suggested to be active on both the plant and the fungal cell wall (Nagy *et al*., 2021). Among the 10 upregulated FCW genes, nine were upregulated in both mutants, while out of the nine down-regulated FCW genes, only three were significantly downregulated in both mutants (**Fig. 4 C and D**; **Table S7**).

### Multiple transcriptional factors are affected by the *cre1* deletion

In total, we found 31 differentially expressed transcription factors in our Quant-Seq data of *C. cinerea cre1* mutants. Of the 31 TFs, 20 TFs were upregulated and 11 TFs were significantly downregulated in at least one of the mutants (**Fig. 5 B; Table S8**). We identified 19 and 23 TFs significantly deregulated in Δ*cre1*-43 and Δ*cre1*-91, respectively. Eleven differentially expressed TFs were common in both mutants, of which nine were upregulated and two were downregulated (**Fig. 5 A**).

**Fig 5.**
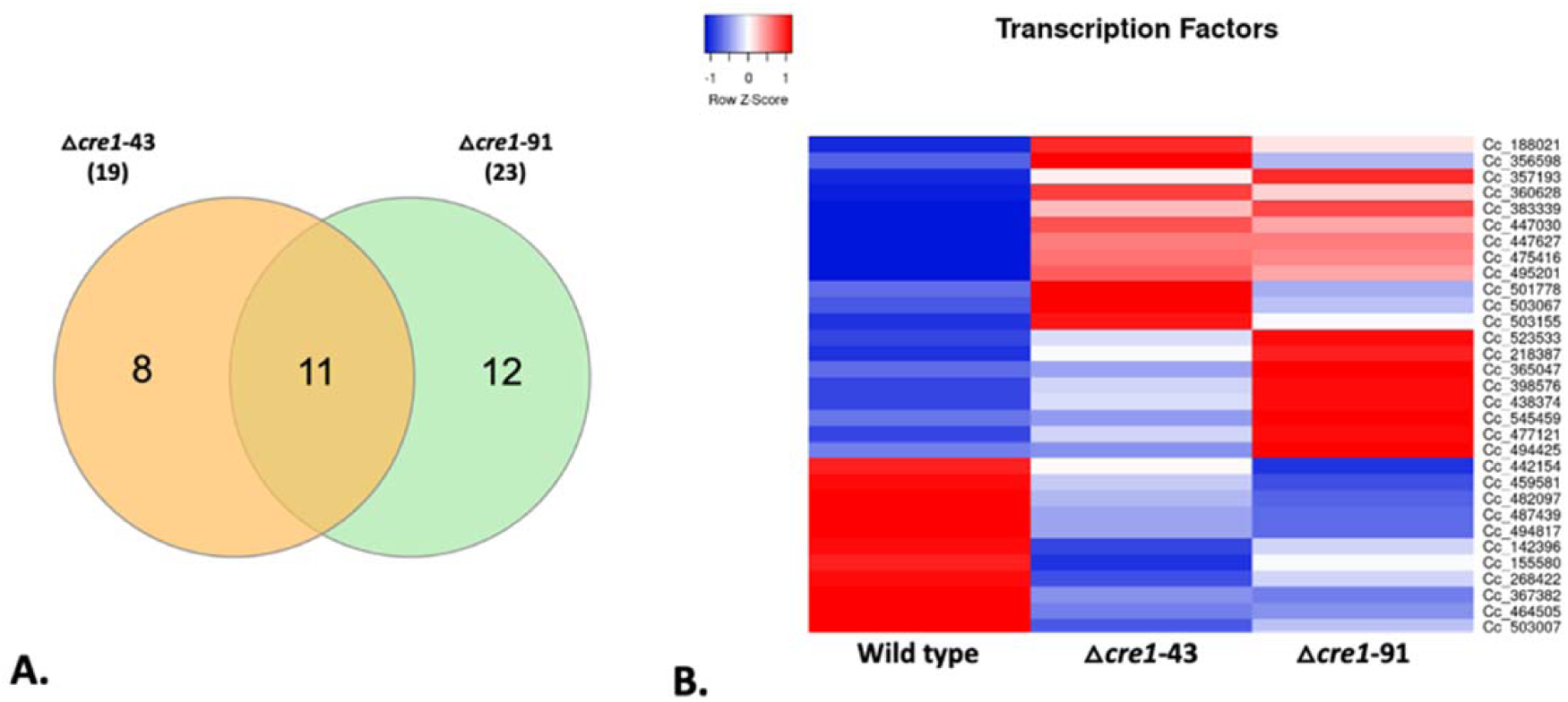
Differentially expressed transcription factor genes in the *C. cinerea* Δ*cre1*-43 and Δ*cre1*-91 mutants compared with wild type strain. (A) A Venn diagram showing the common significantly deregulated genes between Δ*cre1*-43 and Δ*cre1*-91 mutants. (B) Heatmap showing the average CPM of transcription factor genes for the wild type, Δ*cre1*-43 and Δ*cre1*-91 mutants.

Genes encoding High Mobility Group box domain containing proteins (HMG superfamily) were most abundant among the differentially expressed genes. Five TFs containing the HMG box domain were differentially expressed, of which three were upregulated and two were downregulated. We also found that the *C. cinerea* ortholog (protein ID: 447627) of the *S. commune roc1* transcription factor was significantly upregulated in both mutants. Roc1 was recently described as a regulator of cellulase genes in *S. commune* (Marian *et al*., 2021). Similarly, a heat shock factor-type gene (protein ID: 523533) related to the *S. cerevisiae skn7* transcription factor was upregulated in both mutants. In addition, the upregulation of an *Ecd* family gene (protein ID: 438374) that shares homology with the *S. pombe sgt1* and a DNA2/NAM7 helicase-like domain containing transcription factor gene (protein ID: 447030) was observed in the data. The full list for the transcription factors is given in **Table S8**.

### Major facilitator superfamily and oligonucleotide transporters are extensively deregulated

In Quant-Seq data from *C. cinerea cre1* mutants, 54 genes encoding plasma membrane-associated transporter proteins showed a significant change in expression (**Fig. 6 A-B**). Twenty-five and 29 of these genes were up and downregulated, respectively (**Table S9**). The most frequently differentially expressed family among these genes was the Major Facilitator Superfamily (20 genes).

**Fig. 6.**
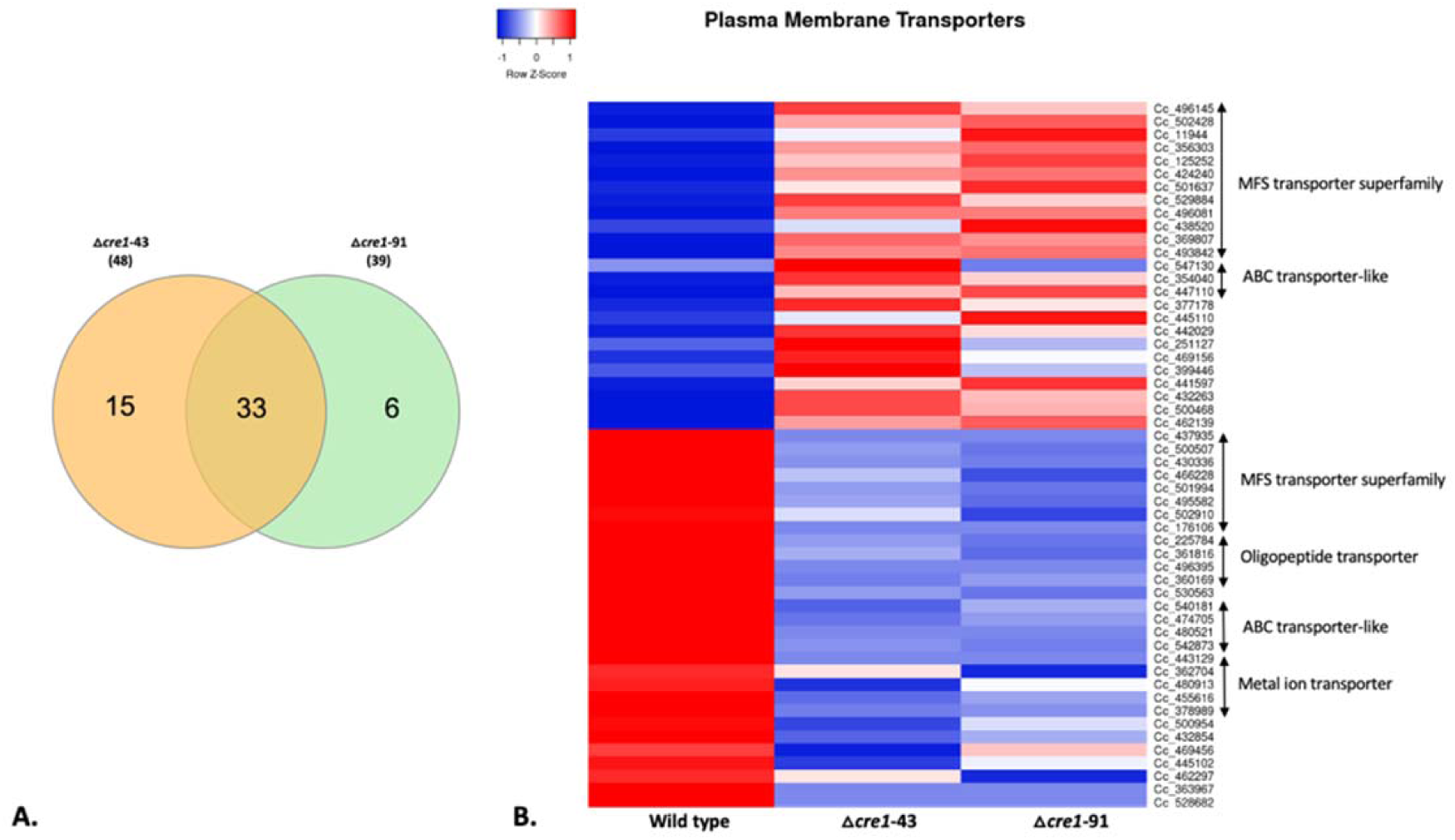
Differentially expressed plasma membrane transporter genes in the *C. cinerea* Δ*cre1*-43 and Δ*cre1*-91 mutants compared with wild type strain. (A) A Venn diagram showing the common significantly affected plasma membrane transporter genes between Δ*cre1*-43 and Δ*cre1*-91 mutants. (B) The heatmap showing the mean CPM across replicates of differentially expressed plasma membrane transporter genes for wild type, Δ*cre1*-43 and Δ*cre1*-91 mutants.

In the Quant-Seq data, four genes containing InterPro domains related to the oligopeptide transporter superfamily were observed to be downregulated in *C. cinerea cre1* mutants. In addition, several ion transporter-related proteins such as phosphate, nickel/cobalt, potassium, chromate, acetate, and zinc/iron permease genes were downregulated (**Table S9**). In addition, several ABC transporter-like proteins, EamA domain, amino acid/polyamine transporter I, purine/cytosine permease, SH3-like domain superfamily, RTA-like protein, and STAS domain superfamily-related genes were also affected, the full list is given in **Table S9**.

## Discussion

In this study, we used a split-marker DNA repair cassette along with Cas9-RNPs to generate knockout strains of the carbon catabolite repressor *cre1* in the model mushroom *Coprinopsis cinerea* and then assessed the transcriptional changes in the mutant strains. *C. cinerea* is a widely used model mushroom to study various molecular aspects of the mushroom biology, including fruiting body development, sexual development and sporulation (Kües, 2000; Kües *et al*., 2016; Kuees *et al*., 2015; Muraguchi *et al*., 2015; Pukkila, 2011; Plaza *et al*., 2014). *C. cinerea* is also an efficient degrader of lignocellulosic plant biomass (Floudas *et al.,* 2012; Hibbett *et al*., 2014). It is a litter decomposer fungus adapted to degrade herbaceous plant biomass (Floudas *et al*., 2020).

Genetic transformation protocols exist for the transformation and gene deletion of *C. cinerea* (Dörnte and Kües, 2012, Sugano *et al*., 2017). The laborious cloning of plasmid-based DNA repair cassettes and low HR-mediated genomic integrations is a major obstacle to performing high-throughput gene deletion studies in *C. cinerea*. Split-marker DNA cassettes gene deletion is commonly used for genetic studies in yeasts and filamentous fungi such as *Fusarium oxysporum* (Chung and Lee, 2015, Pareek *et al*., 2019), but has not yet been applied to *C. cinerea*. Here, we used a split-marker DNA transformation approach with or without *in-vitro* assembled Cas9-RNPs. The results show that Cas9-RNPs with split-marker DNA repair cassettes increased the number of colonies more than six times compared to that without Cas9-RNPs. No *cre1* deletion strain was found among the examined transformants generated using the split-maker DNA cassette without Cas9-RNPs. The increase in the number of transformants and the identification of *C. cinerea cre1* deletion strains by using the split-marker DNA cassette with the Cas9-RNPs approach suggests that Cas9-mediated target-specific nicking may increase site-specific homologous recombination, which ultimately contributes to the integration of foreign DNA into the genome. Both Δ*cre1*-43 and Δ*cre1*-91 mutants showed integration of the *pab1* marker cassette into the *C. cinerea cre1* gene locus, demonstrating that the split-marker approach along with Cas9-RNPs is efficient for targeted gene deletion in *C. cinerea*.

Fruiting body formation is not significantly impaired in *C. cinerea cre1* deleted strains compared to wild-type strains. Further studies are needed to understand the role of *C. cinerea cre1* in fruiting body formation on different carbon sources.

We identified 658-682 upregulated and 280-399 downregulated genes in the *cre1* mutants of *C. cinerea*; these showed a large overlap between the two independent mutants. It is important to note that these probably include genes both under direct or indirect control of Cre1 and assays of direct promoter binding (*e.g.,* ChIP-Seq) will be necessary to distinguish direct and indirect regulation. Nevertheless, the presence of both up- and downregulated genes suggests that Cre1 can act both as a repressor and an activator, respectively, consistent with previous reports (Portnoy *et al*; 2011; Antonieto *et al*., 2014; Peng *et al*., 2021). Differentially expressed genes also provide information on the potential regulon size of Cre1 in *C. cinerea,* which seems to be comparable to that reported in other species. In *T. reesei* 815 and 697 genes were modulated by CRE1 in cellulose and glucose, respectively, and Δ*cre1* showed that 905 genes were upregulated or downregulated in at least one of the conditions (Antonieto *et al*., 2014). For a *A. niger* Δ*creA* mutant 1,423 and 3,077 genes, around 12% and 26% of the total encoded genes, were differentially expressed for the wheat bran and sugar beet pulp respectively was detected at the early time point (Peng *et al*., 2021). In *Dichomitus squalens* around 7% (1,042/15,295) genes were repressed in glucose mediated repression (Daly *et al*., 2019).

The upregulated genes of *C. cinerea cre1* mutants were enriched for GO terms related to carbohydrate metabolism (**Table S6**), which was evident also in the analysis of PCWDE gene expression (**Fig 4**). The upregulation of carbohydrate metabolism related genes in *cre1* mutants is a phenotypic confirmation of the deletion of the *cre1* gene in *C. cinerea*. The role of the *cre1* gene in carbon metabolism, particularly in the CCR, is already well-known for ascomycete fungi, but very little information is available for basidiomycetes (Adnan *et al*., 2017; Gancedo, 1998; Dowzer *et al*., 1991; Ries *et al*., 2016). Among the downregulated genes, it is interesting to see that the expression of genes related to GPCR signalling; further studies can be performed to understand this. Previously, in *A. nidulans* it was suggested that trimeric G-proteins have a role in CreA-independent regulation of CCR (Kunitake *et al*., 2019). The downregulation of G protein-related GO terms in the mutants suggests that the *C. cinerea cre1* gene may be responsible for the normal expression of GPCR proteins in basidiomycetes. Oxidoreductase activity is also enriched in the down-regulated genes of *C. cinerea cre1* mutants. Oxidoreductase activity might be related to auxiliary redox enzymes, which include LMEs (Levasseur *et al*., 2013; Drula *et al*., 2022). The GO terms affected in *cre1* gene deletion in various Ascomycetes fungal species are mainly carbohydrate metabolism, hydrolase activity, transporter activity, oxidoreductase activity, membrane and transcriptional regulation (Peng *et al*., 2021; Antonieto *et al*., 2014), similar GO terms were enriched in the *C. cinerea cre1* mutants as well. This indicates that the functional role of the *cre1* gene is conserved across phyla.

Our results indicated a broad upregulation of PCWDE encoding CAZyme genes in the deletion strains even in the presence of glucose, suggesting that they are repressed either directly or indirectly by *C. cinerea* Cre1. This is consistent with the behaviour of Cre1 orthologs in other species (Peng *et al*., 2021; Yoav *et al*., 2018, De Vries and Makela, 2020, Adnan *et al*., 2017), that is, repression of PCWDEs when the preferred carbon source, such as glucose, is present. In *P. ostreatus*, overexpression of *cre1* resulted in an increase in LMEs and a decrease in cellulose-degrading enzymes, whereas *cre1* knockout strains had a lower amount of LMEs and a slightly more carbohydrate-degrading enzymes (Yoav *et al*., 2018). Our results also showed that cellulose-degrading enzyme genes were mostly upregulated, whereas LMEs are downregulated in *C. cinerea cre1* mutants. In the white rot basidiomycete *D. squalens*, glucose was reported to repress genes related to polysaccharide-degrading CAZymes and carbon catabolic genes (Daly *et al*., 2019). Similarly, glucose-induced reduction of cellulolytic enzyme expression was also shown in *P. ostreatus* (Alfaro *et al*., 2020). In our analysis, we show that deletion of *C. cinerea cre1* leads to upregulation of PCWDE CAZymes even in the presence of glucose. This suggests that the *C. cinerea cre1* gene is the main regulator of PCWDE expression during glucose-mediated CCR in the basidiomycete *C. cinerea*. Broadly, it can be concluded that *cre1*-mediated regulation of CAZymes in *C. cinerea* basidiomycetes is similar to that in Ascomycetes.

Though, as a litter decomposer *C. cinerea* has somewhat reduced set of lignin-degrading enzyme genes compared to white rot species (e.g. *P. ostreatus*)(Floudas et al 2020), these genes were mostly downregulated in the *cre1* mutants. This suggests that genes encoding LMEs fall under a different regulation than cellulases, hemicellulases and pectinases and is consistent with a recent analysis of the secretome of Cre1 deletion and overexpression strains in *P. ostreatus* (Yoav *et al*. 2018). The regulation of LMEs in lignocellulose-degrading basidiomycetes appears to be an interesting research avenue.

CreA in ascomycetes is known to exert its repressive effect on PCWDEs both directly and indirectly, through regulating and interacting with other transcription factors (Wu *et al*., 2020, De Vries and Makela, 2020). These transcription factors include ones specifically responsible for the activation of substrate-specific sets of enzyme-encoding PCWDEs, such as xylanases, cellulases, pectinases (Benocci *et al*., 2017, Thieme *et al*., 2017, Klaubauf *et al*., 2014). While several TFs regulating specific PCWDE genes have been described in the Ascomycetes, only a single gene, Roc1 of *S. commune* (Marian *et al*., 2021) has been associated with cellulose degradation in the Basidiomycota. Our data revealed several transcription factors that are differentially expressed in *C. cinerea cre1* mutants, providing candidates for novel TFs related to plant biomass utilisation.

First, the *C. cinerea* ortholog of Roc1 ortholog is upregulated in both independent *C. cinerea cre1* mutants. The *Schizophyllum commune* transcription factor gene *roc1*, which is conserved among Agaricomycetes, was strongly upregulated in the cellulose containing medium and knockouts showed reduced growth on the cellulose medium, demonstrating its importance in regulating cellulose degrading genes (Marian *et al*., 2021). ChIP-Seq analysis of *S. commune* Roc1 identified promoters of genes involved in lignocellulose degradation, particularly LPMOs (Marian *et al*., 2021). The upregulation of the *C. cinerea* Roc1 ortholog in *C. cinerea cre1* mutants may reveal a conserved link between the two transcription factors in regulating lignocellulose degradation.

We found that the five putative transcription factors belonging to the HMG superfamily are deregulated in the *C. cinerea cre1* mutants. The HMG superfamily is a very large and diverse group of chromosome-binding proteins that are mainly involved in transcriptional regulation, chromatin remodelling, DNA repair, replication, and recombination by altering chromatin architecture (Goodwin *et al*., 1973; Yuan *et al*., 2020; Mallik *et al*., 2018). In *S. cerevisiae*, a gene containing an HMG1 box domain encodes the mitochondrial histone protein Abf2p, which can be phosphorylated by the cAMP-dependent protein kinase, a strain with a defective *abf2p* allele has severe effects on the regulation of mitochondrial DNA content during glucose repression (Cho *et al*., 2001). Deregulation of the HMG box superfamily in *C. cinerea cre1* deletion strains suggests that their interaction may control widespread gene regulation during CCR. Moreover, we also found a *C. cinerea* (523533) gene, which is reciprocal best blast hit to *N. crassa rrg-2* and *S. cerevisiae skn7, and* was upregulated in both mutants. *Rrg-2*/*skn7* is a general stress regulatory transcription factor that affects diverse processes (Jones *et al*., 2007; Raitt *et al*., 2000), including ER stress related to lignocellulase secretion (Fan e*t al*., 2015). Whether the upregulation of the *C. cinerea* ortholog of *rrg-2* is the direct consequence of the deletion of *cre1*, or that of ER-stress caused by the upregulation of PCWDEs remains to be investigated; nevertheless, its these observations suggests that ER stress arises during lignocellulase secretion in the basidiomycetes too.

Another widely conserved transcription factor of the *Ecd* family that shares homology with the *S. pombe sgt1* gene is upregulated in the *C. cinerea* Δ*cre1-91* mutant. *S. pombe Sgt1* regulates carbohydrate and amino acid metabolism (Kainou *et al*., 2006). Although this group of transcription factors is functionally poorly known, *C. cinerea cre1*-mediated regulation of the *C. cinerea* ortholog of *sgt1* may be another way of controlling carbohydrate metabolism. Interestingly, DNA2/NAM7 helicase like putative transcription factor is also strongly upregulated in both independent *C. cinerea cre1* mutants. The DNA2/NAM7 helicase-like protein is involved in unwinding DNA or RNA and affects DNA replication and transcription (De la Cruz *et al*., 1999; M. Plank *et al*., 2018; Gloor *et al*., 2012). Truncated *CRE1* (CRE1-96) in *T. reesei* (Rut-C30 strain) has been reported to upregulate expression of helicase-like transcription factors (Mello-de-Sousa *et al*., 2014). This suggests that helicases can be repressed by Cre1 to effectively orchestrate CCR in basidiomycetes. How the upregulation of this gene is linked to the deletion of *cre1* needs further investigation, we speculate it may be an indirect effect, which is spurred by the need for intense transcription in the hyphae.

Previously in Cre1 was reported in many fungal species to regulate the expression of the MFS transporter genes (Daly *et al*., 2019; Peng *et al*., 2021). In *D. squalens*, sugar transmembrane transporter activity was enriched in genes repressed on Avicel supplemented with glucose compared to Avicel alone (Daly *et al*., 2019). In *N. crassa*, Cre-1 has been shown to regulate not only PCWDEs and other TFs, but also sugar transporter genes that mediate the uptake of decomposition intermediates (Wu *et al*., 2020). Indeed, GO enrichment in the *C. cinerea cre1* mutants suggested that several genes related to transport were affected. These mainly included genes belonging to the Major Facilitator Superfamily. It is known that proteins containing major facilitator superfamily and sugar/inositol-like domains are mainly involved in the transport of simple sugars, oligosaccharides, drugs, amino acids, nucleosides, organic alcohols, and acids (Pao *et al*., 1998; Mueckler *et al*., 1985). The deregulation of several transporters in our data suggests that they may be under the control of *C. cinerea* Cre1 to regulate the transport of specific sugars, peptides, and various ions during CCR.

Finally, our results support that the use of pre-assembled Cas9-RNPs together with a split-marker DNA repair cassette is an efficient approach for gene deletion in *C. cinerea*. Our data demonstrate that *C. cinerea cre1* acts as the carbon catabolite repressor in this fungus and its deletion affects the expression of genes related to carbon metabolism, CAZymes especially PCWDEs upregulation, sugar transporters, ion transporters, and several transcription factors in *C. cinerea*. It also supports the notion that Cre1 achieves a tight regulation of CCR through a combined repression of PCWDEs, transcription factors that positively regulate PCWDEs (*e.g.* Roc1) and membrane transporters which could import simple sugars that can induce the expression of PWCDEs (Wu *et al*., 2020). In the future, the specific targets of the *C. cinerea cre1* gene can be identified, and the role of various transcription factors, transporters can be studied in more detail to understand the CCR and carbon metabolism in wood-degrading fungi.

## Acknowledgment

This work was funded by the Momentum Program of the Hungarian Academy of Sciences (LP2019-13/2019) and by the European Research Council (Grant No. 758161) (both to LGN).

## Supporting Information

**Fig. S1.**
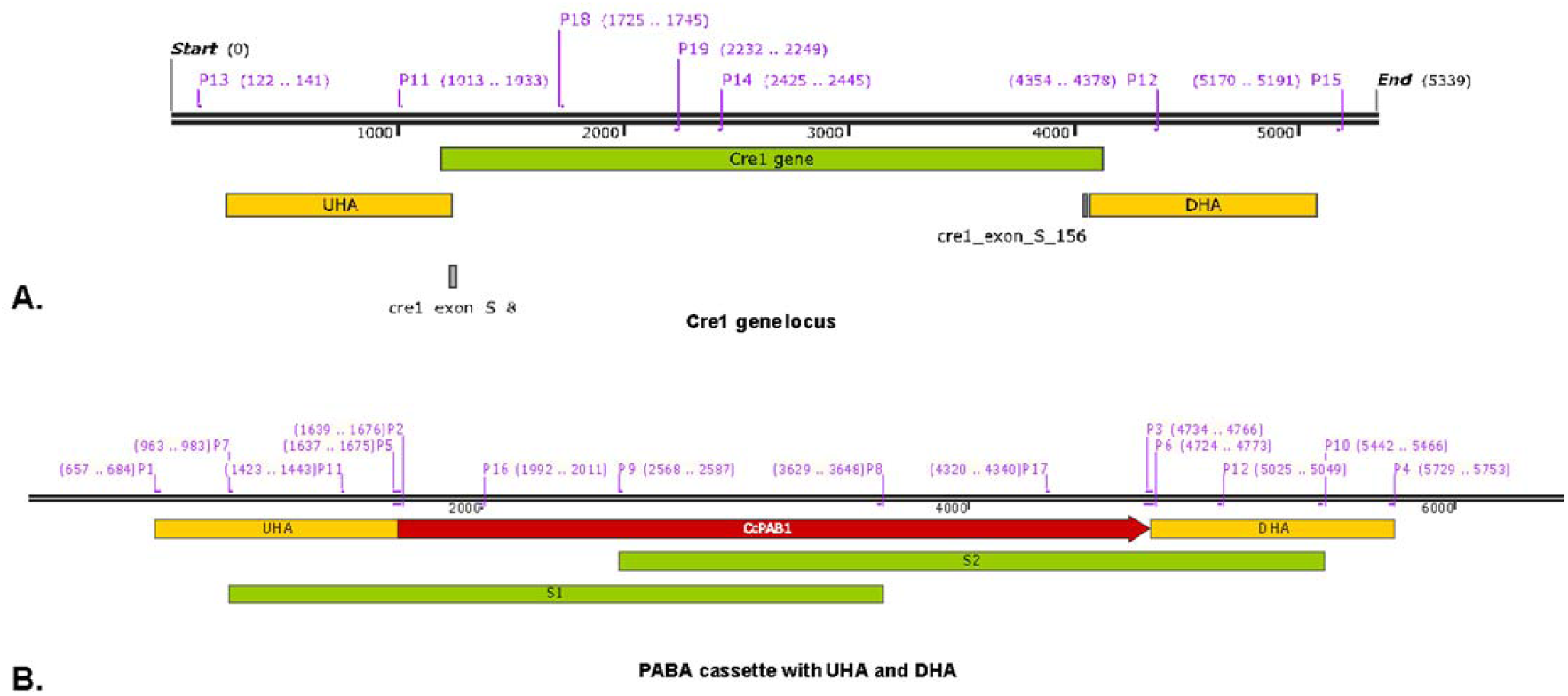
(**A**) DNA map shows the *C. cinerea cre1* gene locus together with the upstream homology arm (UHA), the downstream homology arm (DHA) and the sgRNA binding sites (cre1_exon_S_8 and cre1_exon_S_156). P13, P11, P18, P19, P14, P12 and P15 are primers used for screening the transformants. (**B**) DNA map of *cre1* PABA cassette showing the position of the selection marker *pab1* gene (CcPAB1 or PABA) along with the upstream homology arm (UHA) and downstream homology arm (DHA). The positions of the split marker cassettes are also indicated as split 1 (S1) and split 2 (S2). The different primer positions used for split cassette preparation, screening, and sequencing of the mutants are shown here (P1, P7, P11, P5, P2, P16, P9, P8, P17, P3, P6, P12, P10, and P4).

**Fig. S2:**
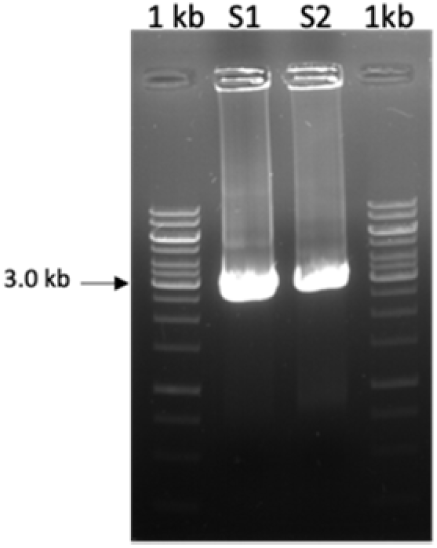
PCR-amplified and purified split 1 (S1) and split 2 (S2) DNA repair cassette used for PEG -mediated protoplast transformation.

**Fig. S3.**
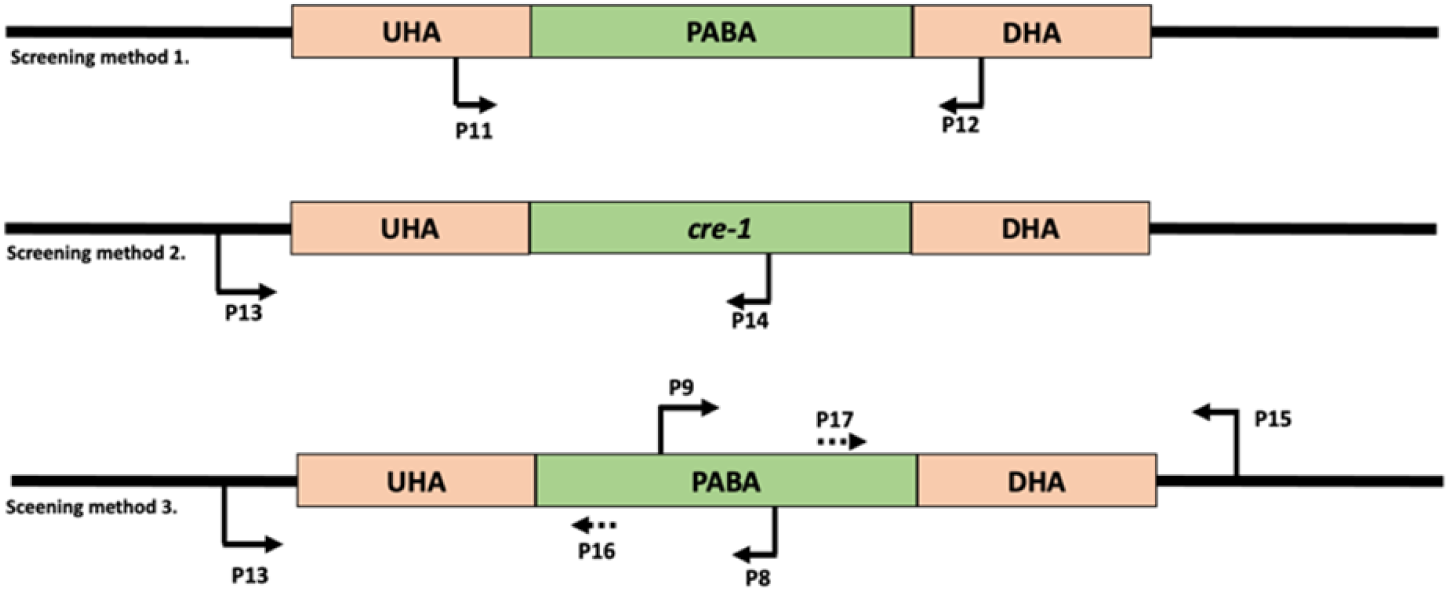
Different screening methods were used to screen the putative *C. cinerea cre1* deleted strain. **Screening method 1**. transformants with PABA integration and wild-type gene PCR amplicons can be distinguished by size using the flanking diagnostic primers (P11 and P12). Here, the size of the wild-type C. cinerea cre1 gene locus = 3.36 KB; the size of the mutant = 3.62 (with PABA insertion). **Screening method 2**. true deletion mutants yield no amplification with the external forward primer P13 and the *cre1* gene internal reverse primer P14. **Screening method 3**. Provide information on the location of the deleted locus using external primer P13 and another PABA internal primer P8. Similarly, at the other end, use primers P9 and P15. P16 and P17 are used for DNA sequencing along with P13 and P15 for final confirmation of the amplified product.

**Fig. S4.**
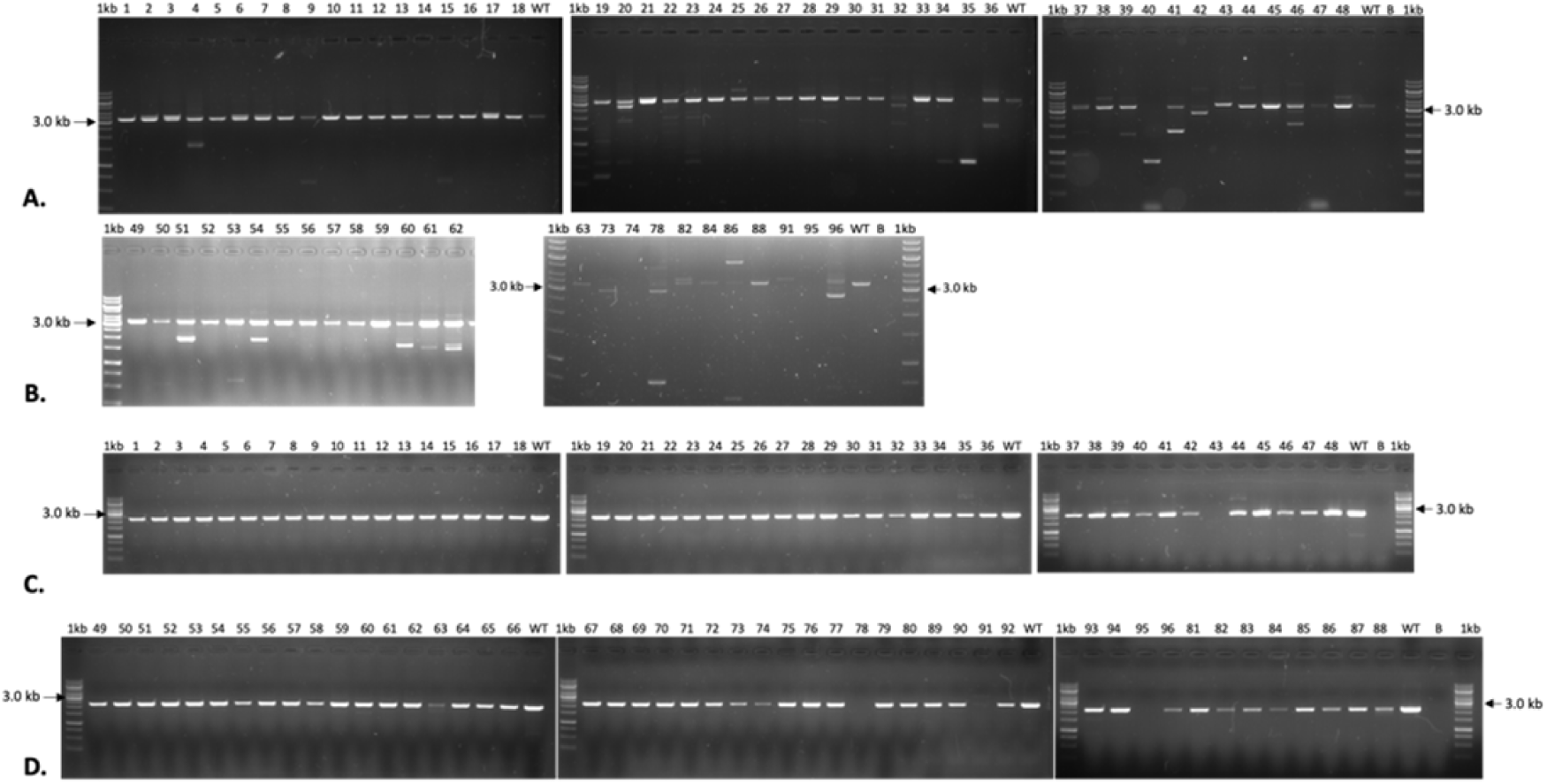
**(A-B)**. PCR screening of the putative *C. cinerea* cre1 mutants (1-96) using screening method 1 (P11 and P12 primers). Colony numbers 1-24; 49-72 were obtained by the transformation approach with split-marker DNA repair cassette alone. Colony numbers 25-48; 73-96 were transformed by the RNPs along with the split marker DNA repair cassette. Not all samples are shown after transformant 63. (**C-D**). PCR screening of *C. cinerea cre1* mutants (1-96) using screening method 2 (P13, P14 primers). The loading order of the samples is discontinuous after transformant 80. WT - Wild type strain, B - blank/water control, 1kb- GeneRuler 1 kb DNA ladder.

**Table S1:** DNA sequences of the sgRNAs protospacer and primers used in the study.

**Table S2:** C. cinerea protein IDs along with its InterPro accession number, InterPro description and Gene Ontology (GO) IDs.

**Table S3:** Quant-seq data analysis of Δ*cre1*-43 mutants compared to wild type strain to obtain log fold change (logFC), average expression and differential expression status (deg status) for C. cinerea protein IDs.

**Table S4:** Analysis of Quant-Seq data of Δcre1-91 mutants compared with the wild-type strain to obtain the log fold change (logFC), average expression and differential expression status (deg status) for the C. cinerea protein IDs.

**Table S5:** Normalized count per million reads (CPM) for Δ*cre1*-43, Δ*cre1*-91, and wild-type samples (Sheet 1). Total number of raw reads, trimmed reads, uniquely mapped reads, and assigned reads obtained from Quant-Seq data for each sample. Also shown is the percentage of uniquely mapped reads compared to the raw reads obtained (Sheet 2).

**Table S6:** The table shows the GO terms along with the corresponding number of genes (annotated, significant) in the up and down-regulated genes of C. cinerea cre1 mutants. Sheet 1: Shows the GO terms in upregulated genes of C. cinerea Δcre1-43 and Δcre1-91 mutants. Sheet 2: Shows the GO terms in down-regulated genes of C. cinerea Δcre1-43 and Δcre1-91 mutants. BP - Biological process, MF - Molecular function and CC - Cellular component.

**Table S7:** Table shows the differentially expressed CAZymes in the *C. cinerea* Δ*cre1*-43 and Δ*cre1*-91 mutants Sheet 1: Shows the PCDWE CAZymes along with their protein IDs, CAZyme classes, plant cell wall components, deg status, and average CPM for the mutants and wild type. Sheet 2: Shows the FCW CAZyme along with their protein IDs, CAZyme classes, fungal cell wall components, deg status and average CPM for the mutants and wild type.

**Table S8:** The differentially expressed transcription factors (protein IDs) in the Δcre1-43 and Δcre1-91 mutants along with their average CPM values, differential expression status (deg status), InterPro description, and the names of their orthologs in other species.

**Table S9:** Putative plasma membrane transporters differentially expressed in Δ*cre1*-43 and Δ*cre1*-91 mutants along with their average CPM values, differential expression status (deg status), InterPro ids, InterPro description and probable location in the cell.

## References

Adnan, M., Zheng, W., Islam, W., Arif, M., Abubakar, Y. S., Wang, Z., & Lu, G. (2017). Carbon catabolite repression in filamentous fungi. International Journal of Molecular Sciences, 19(1), 48.

Alexa A, Rahnenfuhrer J (2021). topGO: Enrichment Analysis for Gene Ontology. R package version 2.46.0. https://bioconductor.org/packages/release/bioc/html/topGO.html.

Alfaro, M., Majcherczyk, A., Kües, U., Ramírez, L., & Pisabarro, A. G. (2020). Glucose counteracts wood-dependent induction of lignocellulolytic enzyme secretion in monokaryon and dikaryon submerged cultures of the white-rot basidiomycete Pleurotus ostreatus. Scientific reports, 10(1), 1–10.

Andrews, S. (2010). FastQC: a quality control tool for high throughput sequence data. https://www.bioinformatics.babraham.ac.uk/projects/fastqc/.

Antonieto, A. C. C., dos Santos Castro, L., Silva-Rocha, R., Persinoti, G. F., & Silva, R. N. (2014). Defining the genome-wide role of CRE1 during carbon catabolite repression in Trichoderma reesei using RNA-Seq analysis. Fungal genetics and biology, 73, 93–103.

Babicki, S., Arndt, D., Marcu, A., Liang, Y., Grant, J. R., Maciejewski, A., & Wishart, D. S. (2016). Heatmapper: web-enabled heat mapping for all. Nucleic acids research, 44(W1), W147–W153.

Benocci, T., Aguilar-Pontes, M. V., Zhou, M., Seiboth, B., & de Vries, R. P. (2017). Regulators of plant biomass degradation in ascomycetous fungi. Biotechnology for biofuels, 10(1), 1–25.

Boontawon, T., Nakazawa, T., Inoue, C., Osakabe, K., Kawauchi, M., Sakamoto, M., & Honda, Y. (2021a). Efficient genome editing with CRISPR/Cas9 in Pleurotus ostreatus. AMB Express, 11(1), 1–11.

Boontawon, T., Nakazawa, T., Xu, H., Kawauchi, M., Sakamoto, M., & Honda, Y. (2021b). Gene targeting using pre-assembled Cas9 ribonucleoprotein and split-marker recombination in Pleurotus ostreatus. FEMS Microbiology Letters.

Bottoli, A. P., Kertesz-Chaloupkova, K., Boulianne, R. P., Aebi, M., & Ku, U. (1999). Rapid isolation of genes from an indexed genomic library of C. cinereus in a novel pab1+ cosmid. Journal of microbiological methods, 35(2), 129–141.

Bushnell, B., Rood, J., & Singer, E. (2017). BBMerge–accurate paired shotgun read merging via overlap. PloS one, 12(10), e0185056.

Capella-Gutiérrez, S., Silla-Martínez, J. M., & Gabaldón, T. (2009). trimAl: a tool for automated alignment trimming in large-scale phylogenetic analyses. Bioinformatics, 25(15), 1972–1973.

Catlett, N. L., Lee, B. N., Yoder, O. C., & Turgeon, B. G. (2003). Split-marker recombination for efficient targeted deletion of fungal genes. Fungal Genet Rep 50: 9–11.

Cherry, J. M., Hong, E. L., Amundsen, C., Balakrishnan, R., Binkley, G., Chan, E. T., et al. (2012). Saccharomyces Genome Database: the genomics resource of budding yeast. Nucleic acids research, 40(D1), D700–D705.

Cho, J. H., Lee, Y. K., & Chae, C. B. (2001). The modulation of the biological activities of mitochondrial histone Abf2p by yeast PKA and its possible role in the regulation of mitochondrial DNA content during glucose repression. Biochimica et Biophysica Acta (BBA)-Gene Structure and Expression, 1522(3), 175–186.

Chung, K. R., & Lee, M. H. (2015). Split-marker-mediated transformation and targeted gene disruption in filamentous fungi. In Genetic Transformation Systems in Fungi, Volume 2 (pp. 175–180). Springer, Cham.

Daly, P., Peng, M., Di Falco, M., Lipzen, A., Wang, M., Ng, V., et al. (2019). Glucose-mediated repression of plant biomass utilization in the white-rot fungus Dichomitus squalens. Applied and environmental microbiology, 85(23), e01828–19.

De Assis, L. J., Silva, L. P., Bayram, O., Dowling, P., Kniemeyer, O., Krüger, T., et al. (2021). Carbon catabolite repression in filamentous fungi is regulated by phosphorylation of the transcription factor CreA. MBio, 12(1), e03146–20

De Jong, J. F., Ohm, R. A., De Bekker, C., Wösten, H. A., & Lugones, L. G. (2010). Inactivation of ku80 in the mushroom-forming fungus Schizophyllum commune increases the relative incidence of homologous recombination. FEMS microbiology letters, 310(1), 91–95.

De la Cruz, J., Kressler, D., & Linder, P. (1999). Unwinding RNA in Saccharomyces cerevisiae: DEAD-box proteins and related families. Trends in biochemical sciences, 24(5), 192–198.

De Vries, R. P., & Mäkelä, M. R. (2020). Genomic and postgenomic diversity of fungal plant biomass degradation approaches. Trends in Microbiology, 28(6), 487–499.

DiCarlo, J. E., Norville, J. E., Mali, P., Rios, X., Aach, J., & Church, G. M. (2013). Genome engineering in Saccharomyces cerevisiae using CRISPR-Cas systems. Nucleic acids research, 41(7), 4336–4343.

Dobin, A., Davis, C. A., Schlesinger, F., Drenkow, J., Zaleski, C., Jha, S., … & Gingeras, T. R. (2013). STAR: ultrafast universal RNA-seq aligner. Bioinformatics, 29(1), 15–21.

Dörnte, B., & Kües, U. (2012). Reliability in transformation of the basidiomycete Coprinopsis cinerea. Current Trends in Biotechnology and Pharmacy, 6(3), 340–355.

Dörnte, B., & Kües, U. (2013). Fast microwave-based DNA extraction from vegetative mycelium and fruiting body tissues of Agaricomycetes for PCR amplification. Current Trends in Biotechnology and Pharmacy, 7(4), 825–836.

Dörnte, B., Peng, C., Fang, Z., Kamran, A., Yulvizar, C., & Kües, U. (2020). Selection markers for transformation of the sequenced reference monokaryon Okayama 7/# 130 and homokaryon AmutBmut of Coprinopsis cinerea. Fungal biology and biotechnology, 7(1), 1–18.

Dowzer, C. E., & Kelly, J. M. (1991). Analysis of the creA gene, a regulator of carbon catabolite repression in Aspergillus nidulans. Molecular and Cellular Biology, 11(11), 5701–5709.

Drula, E., Garron, M. L., Dogan, S., Lombard, V., Henrissat, B., & Terrapon, N. (2022). The carbohydrate-active enzyme database: functions and literature. Nucleic acids research, 50(D1), D571–D577.

Fan, F., Ma, G., Li, J., Liu, Q., Benz, J. P., Tian, C., & Ma, Y. (2015). Genome-wide analysis of the endoplasmic reticulum stress response during lignocellulase production in Neurospora crassa. Biotechnology for biofuels, 8(1), 1–17.

Fasoyin, O. E., Wang, B., Qiu, M., Han, X., Chung, K. R., & Wang, S. (2018). Carbon catabolite repression gene creA regulates morphology, aflatoxin biosynthesis and virulence in Aspergillus flavus. Fungal Genetics and Biology, 115, 41–51

Floudas, D., Bentzer, J., Ahrén, D., Johansson, T., Persson, P., & Tunlid, A. (2020). Uncovering the hidden diversity of litter-decomposition mechanisms in mushroom-forming fungi. The ISME journal, 14(8), 2046–2059.

Floudas, D., Binder, M., Riley, R., Barry, K., Blanchette, R. A., Henrissat, B., et al. (2012). The Paleozoic origin of enzymatic lignin decomposition reconstructed from 31 fungal genomes. Science, 336(6089), 1715–1719.

Floudas, D., Held, B. W., Riley, R., Nagy, L. G., Koehler, G., Ransdell, A. S., et al. (2015). Evolution of novel wood decay mechanisms in Agaricales revealed by the genome sequences of Fistulina hepatica and Cylindrobasidium torrendii. Fungal Genetics and Biology, 76, 78–92.

Gancedo, J. M. (1998). Yeast carbon catabolite repression. Microbiology and molecular biology reviews, 62(2), 334–361.

Gloor, J. W., Balakrishnan, L., Campbell, J. L., & Bambara, R. A. (2012). Biochemical analyses indicate that binding and cleavage specificities define the ordered processing of human Okazaki fragments by Dna2 and FEN1. Nucleic acids research, 40(14), 6774–6786.

Goodwin, G. H., Sanders, C., & Johns, E. W. (1973). A new group of chromatin-associated proteins with a high content of acidic and basic amino acids. European Journal of Biochemistry, 38(1), 14–19.

Görke, B., & Stülke, J. (2008). Carbon catabolite repression in bacteria: many ways to make the most out of nutrients. Nature Reviews Microbiology, 6(8), 613–624.

Grunwald, P. (2016). Handbook of carbohydrate-modifying biocatalysts. Jenny Stanford Publishing.

Heberle, H., Meirelles, G. V., da Silva, F. R., Telles, G. P., & Minghim, R. (2015). InteractiVenn: a web-based tool for the analysis of sets through Venn diagrams. BMC bioinformatics, 16(1), 1–7.

Hibbett, D. S., Bauer, R., Binder, M., Giachini, A. J., Hosaka, K., Justo, A., et al. (2014). 14 Agaricomycetes. In Systematics and evolution (pp. 373–429). Springer, Berlin, Heidelberg.)

Horton, P., Park, K. J., Obayashi, T., & Nakai, K. (2006). Protein subcellular localization prediction with WoLF PSORT. In Proceedings of the 4th Asia-Pacific bioinformatics conference (pp. 39-48). https://doi.org/10.1142/9781860947292_0007

Hu, Y., Xu, W., Hu, S., Lian, L., Zhu, J., Shi, L., et al. (2020). In Ganoderma lucidum, Glsnf1 regulates cellulose degradation by inhibiting GlCreA during the utilization of cellulose. Environmental microbiology, 22(1), 107–121

Ishibashi, K., Suzuki, K., Ando, Y., Takakura, C., & Inoue, H. (2006). Nonhomologous chromosomal integration of foreign DNA is completely dependent on MUS-53 (human Lig4 homolog) in Neurospora. Proceedings of the National Academy of Sciences, 103(40), 14871–14876.

Jones, C. A., Greer-Phillips, S. E., & Borkovich, K. A. (2007). The response regulator RRG-1 functions upstream of a mitogen-activated protein kinase pathway impacting asexual development, female fertility, osmotic stress, and fungicide resistance in Neurospora crassa. Molecular biology of the cell, 18(6), 2123–2136.

Kainou, T., Shinzato, T., Sasaki, K., Mitsui, Y., Giga-Hama, Y., Kumagai, H., & Uemura, H. (2006). Spsgt1, a new essential gene of Schizosaccharomyces pombe, is involved in carbohydrate metabolism. Yeast, 23(1), 35–53.

Klaubauf, S., Narang, H. M., Post, H., Zhou, M., Brunner, K., Mach-Aigner, A. R., et al. (2014). Similar is not the same: differences in the function of the (hemi-) cellulolytic regulator XlnR (Xlr1/Xyr1) in filamentous fungi. Fungal Genetics and Biology, 72, 73–81.

Kohler, A., Kuo, A., Nagy, L. G., Morin, E., Barry, K. W., Buscot, F., et al. (2015). Convergent losses of decay mechanisms and rapid turnover of symbiosis genes in mycorrhizal mutualists. Nature genetics, 47(4), 410–415.

Krappmann, S. (2007). Gene targeting in filamentous fungi: the benefits of impaired repair. Fungal biology reviews, 21(1), 25–29.

Krizsan, K., Almási, É., Merényi, Z., Sahu, N., Virágh, M., Kószó, T., et al. (2019). Transcriptomic atlas of mushroom development reveals conserved genes behind complex multicellularity in fungi. Proceedings of the National Academy of Sciences, 116(15), 7409–7418.

Kück, U., & Hoff, B. (2010). New tools for the genetic manipulation of filamentous fungi. Applied microbiology and biotechnology, 86(1), 51–62.

Kuees, U., & Navarro-Gonzalez, M. (2015). How do Agaricomycetes shape their fruiting bodies? 1. Morphological aspects of development. Fungal Biology Reviews, 29(2), 63–97.

Kües, U. (2000). Life history and developmental processes in the basidiomycete Coprinus cinereus. Microbiology and molecular biology reviews, 64(2), 316–353.

Kües, U., Subba, S., Yu, Y., Sen, M., Khonsuntia, W., Singhaduang, W., et al. (2016). Regulation of fruiting body development in Coprinopsis cinerea. Mushroom Sci, 19, 318–322.

Kunitake, E., Li, Y., Uchida, R., Nohara, T., Asano, K., Hattori, A., et al. (2019). CreA-independent carbon catabolite repression of cellulase genes by trimeric G-protein and protein kinase A in Aspergillus nidulans. Current genetics, 65(4), 941–952

Levasseur, A., Drula, E., Lombard, V., Coutinho, P. M., & Henrissat, B. (2013). Expansion of the enzymatic repertoire of the CAZy database to integrate auxiliary redox enzymes. Biotechnology for biofuels, 6(1), 1–14.

Liao, Y., Smyth, G. K., & Shi, W. (2014). featureCounts: an efficient general purpose program for assigning sequence reads to genomic features. Bioinformatics, 30(7), 923–930.

Liu, K., Sun, B., You, H., Tu, J. L., Yu, X., Zhao, P., & Xu, J. W. (2020). Dual sgRNA-directed gene deletion in basidiomycete Ganoderma lucidum using the CRISPR/Cas9 system. Microbial biotechnology, 13(2), 386–396.

Lou, H., Ye, Z., Yun, F., Lin, J., Guo, L., Chen, B., & Mu, Z. (2018). Targeted gene deletion in Cordyceps militaris using the split-marker approach. Molecular biotechnology, 60(5), 380–385.

Löytynoja, A. (2014). Phylogeny-aware alignment with PRANK. In Multiple sequence alignment methods (pp. 155–170). Humana Press, Totowa, NJ.

M. Plank, T. D., & Wilkinson, M. F. (2018). RNA decay factor UPF1 promotes protein decay: a hidden talent. BioEssays, 40(1), 1700170.

Makinen, M., Kuuskeri, J., Laine, P., Smolander, O. P., Kovalchuk, A., Zeng, Z., et al. (2019). Genome description of Phlebia radiata 79 with comparative genomics analysis on lignocellulose decomposition machinery of phlebioid fungi. BMC genomics, 20(1), 1–22.

Mallik, R., Kundu, A., & Chaudhuri, S. (2018). High mobility group proteins: the multifaceted regulators of chromatin dynamics. The Nucleus, 61(3), 213–226.

Marian, I. M., Vonk, P. J., Valdes, I. D., Barry, K., Bostock, B., Carver, A., et al. (2021). The transcription factor Roc1 is a regulator of cellulose degradation in the wood-decaying mushroom Schizophyllum commune. bioRxiv.

Mello-de-Sousa, T. M., Gorsche, R., Rassinger, A., Poças-Fonseca, M. J., Mach, R. L., & Mach-Aigner, A. R. (2014). A truncated form of the carbon catabolite repressor 1 increases cellulase production in Trichoderma reesei. Biotechnology for biofuels, 7, 1–12.

Miyauchi, S., Kiss, E., Kuo, A., Drula, E., Kohler, A., & Sánchez-García, M. et al. (2020). Large-scale genome sequencing of mycorrhizal fungi provides insights into the early evolution of symbiotic traits. Nature Communications, 11(1). doi: 10.1038/s41467-020-18795-w

Miyauchi, S., Navarro, D., Grisel, S., Chevret, D., Berrin, J. G., & Rosso, M. N. (2017). The integrative omics of white-rot fungus Pycnoporus coccineus reveals co-regulated CAZymes for orchestrated lignocellulose breakdown. PloS one, 12(4), e0175528.

Mueckler, M., Caruso, C., Baldwin, S. A., Panico, M., Blench, I., Morris, H. R., et al. (1985). Sequence and structure of a human glucose transporter. Science, 229(4717), 941–945.

Muraguchi, H., Umezawa, K., Niikura, M., Yoshida, M., Kozaki, T., Ishii, K., et al. (2015). Strand-specific RNA-seq analyses of fruiting body development in Coprinopsis cinerea. PloS one, 10(10), e0141586.

Murphy, C., Powlowski, J., Wu, M., Butler, G., & Tsang, A. (2011). Curation of characterized glycoside hydrolases of fungal origin. Database, 2011.

Nagy, L. G., Riley, R., Tritt, A., Adam, C., Daum, C., Floudas, D., et al. (2016). Comparative genomics of early-diverging mushroom-forming fungi provides insights into the origins of lignocellulose decay capabilities. Molecular biology and evolution, 33(4), 959–970.

Nagy, L. G., Vonk, P. J., Kunzler, M., Foldi, C., Viragh, M., Ohm, R. A., et al. (2021). Lessons on fruiting body morphogenesis from genomes and transcriptomes of Agaricomycetes. bioRxiv.

Nakazawa, T., Ando, Y., Kitaaki, K., Nakahori, K., & Kamada, T. (2011). Efficient gene targeting in ΔCc. ku70 or ΔCc. lig4 mutants of the agaricomycete Coprinopsis cinerea. Fungal Genetics and Biology, 48(10), 939–946.

Ninomiya, Y., Suzuki, K., Ishii, C., & Inoue, H. (2004). Highly efficient gene replacements in Neurospora strains deficient for nonhomologous end-joining. Proceedings of the National Academy of Sciences, 101(33), 12248–12253.

Ohm, R. A., de Jong, J. F., Berends, E., Wang, F., Wösten, H. A., & Lugones, L. G. (2010). An efficient gene deletion procedure for the mushroom-forming basidiomycete Schizophyllum commune. World journal of microbiology and biotechnology, 26(10), 1919–1923.

Pao, S. S., Paulsen, I. T., & Saier Jr, M. H. (1998). Major facilitator superfamily. Microbiology and molecular biology reviews, 62(1), 1–34.

Pareek, M., Almog, Y., Bari, V. K., Hazkani-Covo, E., Onn, I., & Covo, S. (2019). Alternative functional rad21 paralogs in Fusarium oxysporum. Frontiers in microbiology, 10, 1370.

Peng, M., Khosravi, C., Lubbers, R. J., Kun, R. S., Pontes, M. V. A., Battaglia, E., et al. (2021). CreA-mediated repression of gene expression occurs at low monosaccharide levels during fungal plant biomass conversion in a time and substrate dependent manner. The Cell Surface, 7, 100050.

Plaza, D. F., Lin, C. W., van der Velden, N. S. J., Aebi, M., & Künzler, M. (2014). Comparative transcriptomics of the model mushroom Coprinopsis cinerea reveals tissue-specific armories and a conserved circuitry for sexual development. BMC genomics, 15(1), 1–17.

Portnoy, T., Margeot, A., Linke, R., Atanasova, L., Fekete, E., Sándor, E., et al. (2011). The CRE1 carbon catabolite repressor of the fungus Trichoderma reesei: a master regulator of carbon assimilation. BMC genomics, 12(1), 1–12.

Pukkila, P. J. (2011). Coprinopsis cinerea. Current Biology, 21(16), R616–R617.

Raitt, D. C., Johnson, A. L., Erkine, A. M., Makino, K., Morgan, B., Gross, D. S., & Johnston, L. H. (2000). The Skn7 response regulator of Saccharomyces cerevisiae interacts with Hsf1 in vivo and is required for the induction of heat shock genes by oxidative stress. Molecular biology of the cell, 11(7), 2335–2347.

Rambaut, 2018 FigTree V1.4.4 https://github.com/rambaut/figtree/releases.

Raschmanová, H., Weninger, A., Glieder, A., Kovar, K., & Vogl, T. (2018). Implementing CRISPR-Cas technologies in conventional and non-conventional yeasts: current state and future prospects. Biotechnology advances, 36(3), 641–665.

Ries, L. N., Beattie, S. R., Espeso, E. A., Cramer, R. A., & Goldman, G. H. (2016). Diverse regulation of the CreA carbon catabolite repressor in Aspergillus nidulans. Genetics, 203(1), 335–352.

Ritchie, M. E., Phipson, B., Wu, D. I., Hu, Y., Law, C. W., Shi, W., & Smyth, G. K. (2015). limma powers differential expression analyses for RNA-sequencing and microarray studies. Nucleic acids research, 43(7), e47–e47.

Robinson, M. D., McCarthy, D. J., & Smyth, G. K. (2010). edgeR: a Bioconductor package for differential expression analysis of digital gene expression data. Bioinformatics, 26(1), 139–140.

Ruiz-Dueñas, F. J., Barrasa, J. M., Sánchez-García, M., Camarero, S., Miyauchi, S., Serrano, A., et al. (2021). Genomic analysis enlightens Agaricales lifestyle evolution and increasing peroxidase diversity. Molecular biology and evolution, 38(4), 1428–1446.

Rytioja, J., Hildén, K., Yuzon, J., Hatakka, A., De Vries, R. P., & Mäkelä, M. R. (2014). Plant-polysaccharide-degrading enzymes from basidiomycetes. Microbiology and Molecular Biology Reviews, 78(4), 614–649.

Sahu, N., Merényi, Z., Bálint, B., Kiss, B., Sipos, G., Owens, R. A., & Nagy, L. G. (2021). Hallmarks of Basidiomycete soft-and white-rot in wood-decay-Omics data of two Armillaria species. Microorganisms, 9(1), 149

Salame, T. M., Knop, D., Tal, D., Levinson, D., Yarden, O., & Hadar, Y. (2012). Predominance of a versatile-peroxidase-encoding gene, mnp4, as demonstrated by gene replacement via a gene targeting system for Pleurotus ostreatus. Applied and environmental microbiology, 78(15), 5341–5352.

Santangelo, G. M. (2006). Glucose signaling in Saccharomyces cerevisiae. Microbiology and Molecular Biology Reviews, 70(1), 253–282.

Schmieder, S. S., Stanley, C. E., Rzepiela, A., van Swaay, D., Sabotič, J., Nørrelykke, S. F., et al. (2019). Bidirectional propagation of signals and nutrients in fungal networks via specialized hyphae. Current Biology, 29(2), 217–228.

Schuster, M., & Kahmann, R. (2019). CRISPR-Cas9 genome editing approaches in filamentous fungi and oomycetes. Fungal Genetics and Biology, 130, 43–53.

Stanley, C. E., Stöckli, M., van Swaay, D., Sabotič, J., Kallio, P. T., Künzler, M., et al. (2014). Probing bacterial–fungal interactions at the single cell level. Integrative Biology, 6(10), 935–945.

Strasser, K., McDonnell, E., Nyaga, C., Wu, M., Wu, S., Almeida, H., et al. (2015). mycoCLAP, the database for characterized lignocellulose-active proteins of fungal origin: resource and text mining curation support. Database, 2015.

Strauss, J., Horvath, H. K., Abdallah, B. M., Kindermann, J., Mach, R. L., & Kubicek, C. P. (1999). The function of CreA, the carbon catabolite repressor of Aspergillus nidulans, is regulated at the transcriptional and post-transcriptional level. Molecular microbiology, 32(1), 169–178.

Sugano, S. S., Suzuki, H., Shimokita, E., Chiba, H., Noji, S., Osakabe, Y., & Osakabe, K. (2017). Genome editing in the mushroom-forming basidiomycete Coprinopsis cinerea, optimized by a high-throughput transformation system. Scientific reports, 7(1), 1–9.

Sun, J., & Glass, N. L. (2011). Identification of the CRE-1 cellulolytic regulon in Neurospora crassa. PLoS One, 6(9), e25654.

Suzuki, H., Igarashi, K., & Samejima, M. (2008). Real-time quantitative analysis of carbon catabolite derepression of cellulolytic genes expressed in the basidiomycete Phanerochaete chrysosporium. Applied microbiology and biotechnology, 80(1), 99–106.

Swamy, S., Uno, I., & Ishikawa, T. (1984). Morphogenetic effects of mutations at the A and B incompatibility factors in Coprinus cinereus. Microbiology, 130(12), 3219–3224.

Thieme, N., Wu, V. W., Dietschmann, A., Salamov, A. A., Wang, M., Johnson, et al. (2017). The transcription factor PDR-1 is a multi-functional regulator and key component of pectin deconstruction and catabolism in Neurospora crassa. Biotechnology for biofuels, 10(1), 1–21.

Vonk, P. J., Escobar, N., Wösten, H. A., Lugones, L. G., & Ohm, R. A. (2019). High-throughput targeted gene deletion in the model mushroom Schizophyllum commune using pre-assembled Cas9 ribonucleoproteins. Scientific reports, 9(1), 1–8.

Wang, Q., & Coleman, J. J. (2019). CRISPR/Cas9-mediated endogenous gene tagging in Fusarium oxysporum. Fungal Genetics and Biology, 126, 17–24.

Wu, V. W., Thieme, N., Huberman, L. B., Dietschmann, A., Kowbel, D. J., Lee, J., et al. (2020). The regulatory and transcriptional landscape associated with carbon utilization in a filamentous fungus. Proceedings of the National Academy of Sciences, 117(11), 6003–6013.

Xie, S., Shen, B., Zhang, C., Huang, X., & Zhang, Y. (2014). sgRNAcas9: a software package for designing CRISPR sgRNA and evaluating potential off-target cleavage sites. PloS one, 9(6), e100448.

Xie, Y., Chang, J., & Kwan, H. S. (2020). Carbon metabolism and transcriptome in developmental paths differentiation of a homokaryotic Coprinopsis cinerea strain. Fungal Genetics and Biology, 143, 103432.

Yoav, S., Salame, T. M., Feldman, D., Levinson, D., Ioelovich, M., Morag, E., et al. (2018). Effects of cre1 modification in the white-rot fungus Pleurotus ostreatus PC9: altering substrate preference during biological pretreatment. Biotechnology for biofuels, 11(1), 1–16.

Yu, J. H., Hamari, Z., Han, K. H., Seo, J. A., Reyes-Domínguez, Y., & Scazzocchio, C. (2004). Double-joint PCR: a PCR-based molecular tool for gene manipulations in filamentous fungi. Fungal genetics and biology, 41(11), 973–981.

Yuan, S., Liu, Z., Xu, Z., Liu, J., & Zhang, J. (2020). High mobility group box 1 (HMGB1): a pivotal regulator of hematopoietic malignancies. Journal of Hematology & Oncology, 13(1), 1–19.

Zhang, D., Yang, Y., Castlebury, L. A., & Cerniglia, C. E. (1996). A method for the large scale isolation of high transformation efficiency fungal genomic DNA. FEMS microbiology letters, 145(2), 261–265.

Zhang, J., Markillie, L. M., Mitchell, H. D., Gaffrey, M. J., Orr, G., & Schilling, J. S. (2022). Distinctive carbon repression effects in the carbohydrate-selective wood decay fungus Rhodonia placenta. Fungal Genetics and Biology, 159, 103673.

